# The protected physiological status of intracellular *Salmonella enterica* persisters reduces host cell-imposed stress

**DOI:** 10.1101/2020.09.22.308114

**Authors:** Marc Schulte, Katharina Olschewski, Michael Hensel

## Abstract

Today, we are faced with increasingly occurring bacterial infections that are hard to treat and often tend to relapse. These recurrent infections can occur possibly due to antibiotic-tolerant persister cells. Antibiotic persistent bacteria represent a small part of a bacterial population that enters a non-replicating (NR) state arising from phenotypic switching. Intracellularly, after uptake by phagocytic cells, *Salmonella enterica* serovar Typhimurium (STM) forms persister cells that are able to subvert immune defenses of the host. However, the clear physiological state and perceptual properties are still poorly understood and many questions remain unanswered. Here we describe further development of fluorescent protein-based reporter plasmids that were used to detect intracellular NR persister cells and monitor the expression of stress response genes via extensive flow cytometric analyses. Moreover, we performed extensive measurements of the metabolic properties of NR STM at the early course of infection. Our studies demonstrate that NR STM persister cells perceive their environment and are capable respond to stress factors. Since persisters showed a lower stress response compared to replicating (R) STM, which was not a consequence of a lower metabolic capacity, the persistent status of STM serves as protective niche. Furthermore, up to 95% of NR STM were metabolically active at the beginning of infection additionally showing no difference in the metabolic capacity compared to R STM. The accessory capability of NR STM persisters to sense and to react to stress with constant metabolic activity may supports the pathogen to create a more permissive environment for recurrent infections.

## Introduction

The global increase of multi-resistant bacteria presents one of the major challenges to human health in the near future. In addition, physicians frequently encounter bacterial infections that are very difficult to treat, and often relapse without the presence of genetic resistance to antibiotics (Blango and Mulvey, 2010; Caygill et al., 1994; Levine et al., 1982; Lin and Flynn, 2010; Mulvey et al., 2001). These recurrent infections can only be defeated by several rounds of antibiotic treatment, possibly due to the presence of antibiotic-tolerant persister cells. In the context of recurrent infections, the heterogeneous phenomenon of bacterial antibiotic persistence is becoming increasingly important. Antibiotic persistence describes a phenomenon in which a small part of a bacterial population enters a non-replicating (NR) state that can survive actually lethal concentrations of antibiotics during infection. Persister cells arise from a genetically clonal bacterial population by a transient and reversible phenotype switch, leading to a NR and multidrug-tolerant subpopulation (Bigger, 1944; reviewed in Harms et al., 2016; Keren et al., 2004).

Recent *in vitro* investigations of antibiotic persistence observed bacteria entering a dormant state when grown in laboratory media (Conlon et al., 2016; Maisonneuve and Gerdes, 2014; Shan et al., 2017). However, the physiological state and the perceptual properties of intracellular persistent bacteria are still poorly understood. Many questions regarding the interface of antibiotic persisters with their host remain to be answered, and there is demand for sensitive tools to interrogate the interplay of both organisms. Better understanding of the physiology of persisters will help to device new forms of antimicrobial therapy.

The facultative intracellular pathogen *Salmonella enterica* causes acute and chronic infections (Gunn et al., 2014), and *S. enterica* serovar Typhimurium (STM) serves as a model organism for a facultative intracellular pathogen forming persister cells during systemic infections. It was shown that intracellular persister cells of STM do not show complete dormancy, but consist of subpopulations of metabolically active bacteria, as well as inactive cells showing decreasing responsiveness to external stimuli over the course of infection (Helaine et al., 2010). Similar NR but metabolically active bacteria have also been observed in macrophages infected with *Mycobacterium tuberculosis* (Manina et al., 2015). It was shown that many persisters of STM are formed immediately upon phagocytosis by macrophages. Vacuolar acidification as well as nutritional deprivation was mentioned as one of the main factors leading to macrophage-induced persister formation (Helaine et al., 2014). More recently, Stapels et al. (2018) reported that persisters of STM translocate SPI2-T3SS effector proteins to dampen proinflammatory innate immune response, and induce anti-inflammatory macrophage polarization. Such reprogramming of their host cells allows NR STM to survive, and might lead to an advantage during infection relapse after termination of antibiosis (Stapels et al., 2018).

Within phagocytic cells, STM encounters harsh environmental conditions and various defense mechanisms including antimicrobial peptides and the respiratory burst (Kagaya et al., 1989; Slauch, 2011; Zhang and Gallo, 2016). For the pathogen it is of crucial importance to sense and react to potentially detrimental factors. Hence, STM has evolved a plethora of defensive virulence mechanisms to withstand antimicrobial effectors and to overcome the clearance by the host (reviewed in Fang et al., 2016). These stress response systems (SRS) are able to sense harmful conditions, as well as perturbations of the bacterial envelope, in periplasm, or in cytoplasm. SRS have to perform efficiently in space and time for successful survival of STM within hazardous host environments.

Despite being equipped with an extensive set of defensive and offensive virulence factors, the individual fate of intracellular STM is highly diverse (Helaine and Holden, 2013). It was shown that stress response and formation of persister cells are closely related. Among other, main mediators for persister formation are considered to be SOS response via RecA and LexA, stringent response via (p)ppGpp, oxidative stress response via OxyR and SoxSR, and toxin-antitoxin modules (reviewed in Gollan et al., 2019; Harms et al., 2016; Prax and Bertram, 2014). But what happens after establishment of the persistent status? Is switch from persister state to normal growth a merely stochastic event, or can persister monitor their environment with the ability to respond to cues? Addressing these questions is challenging due to population heterogeneity of intracellular STM, the low frequency of persisters, and their low metabolic activity.

Perception of environmental stimuli and especially stressors are of crucial importance for pathogens to maintain their cell integrity. So far, little is known about the stress response of intracellular NR STM. We investigated whether NR STM after entering persistence are still capable in sensing stress factors and in responding by inducing stress responses. If so, is the level of stress response of NR STM similar to replicating (R) STM, or altered? To address these questions, we developed further the recently introduced reporter system (Schulte et al., 2020) to enable analyses of stress response of intracellular STM persister cells. Here we introduce dual fluorescence reporters which enable extremely sensitive flow cytometric analyses of both, stress response and metabolic properties of intracellular STM persisters. Our analyses reveal that intracellular NR STM persister cells perceive their environment and respond to stressors. NR and R STM can mount responses to stressors, and the low response observed for NR STM indicates that the persistent status of intracellular STM serves as protective niche.

## Results

### Design of fluorescence protein-based reporters for stress response of non-replicating intracellular Salmonella enterica at single cell level

In this study, we analyzed the stress response of NR, persisting intracellular STM at the single cell level. For this purpose, we changed the basic design of dual fluorescence stress reporters (Schulte et al., 2020) as shown in **Fig. 1**. The constitutive EM7 promoter was replaced by the tet-ON cassette (Schulte et al., 2019) to induce expression of a gene of interest (GOI) by addition of the non-antibiotic inducer anhydrotetracycline (AHT). The resulting dual fluorescence reporters consisted of the tet-ON cassette for controlled expression of DsRed version DsRed T3_S4T (Sorensen et al., 2003), and sfGFP under control of the regulated promoters of SRS genes *msrA, trxA,* or *htrA* (**Fig. 1A**).

**Fig. 1:**
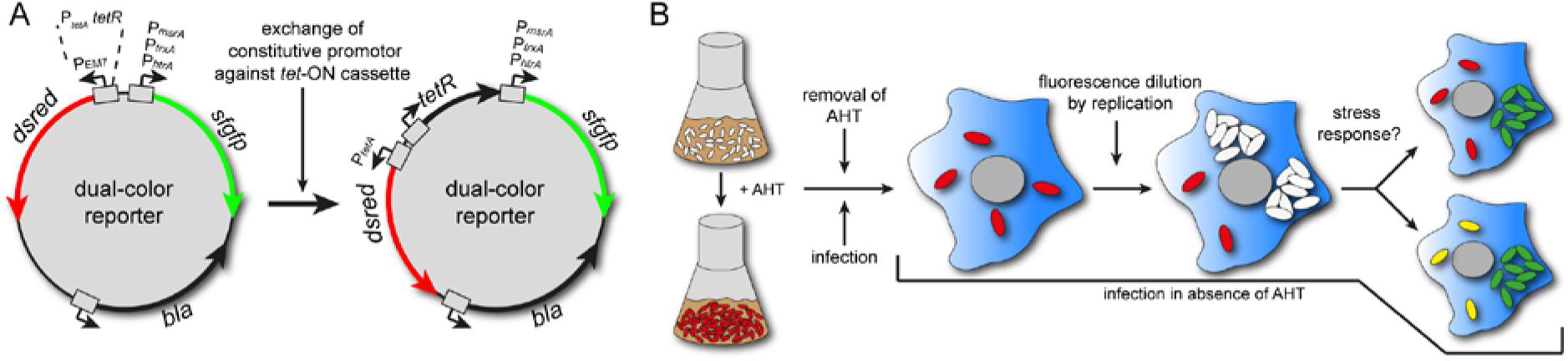
Design of reporters to measure stress response of non-replicating STM. A) For detection of NR intracellular STM, the EM7 promoter of dual FP reporters for *msrA*, *trxA,* or *htrA* was replaced by the tet-ON cassette to enable controlled induction of *dsred* expression by AHT. B) Addition of AHT to growing cultures of STM induced DsRed synthesis. Before infection of macrophages, AHT was removed by centrifugation and washing. Infection was performed in absence of AHT. After infection, R intracellular STM dilute cellular DsRed levels. NR intracellular STM maintain cellular DsRed levels and were detected as DsRed-positive events. In addition, the *msrA*-induced sfGFP intensity provides information about response to stress imposed by the host cell.

The principle for the detection of persisters was adapted from Helaine et al. (2010). Addition of AHT to subcultures used as infection inoculum resulted in synthesis of DsRed. Before infection, AHT was removed by centrifugation and washing to terminate further DsRed synthesis. The principle of fluorescence protein (FP) dilution is that intracellular R STM continuously dilute DsRed, thus, decrease fluorescence levels, while NR or persistent STM maintain DsRed and consistent fluorescence levels. Furthermore, the sfGFP signal reported exposure to stressors and induction of SRS (**Fig. 1B**).

To confirm function of newly generated reporter plasmids, we used *in vitro* cultures of STM WT harboring reporter [P*_tetA_*::*dsred* P*_msrA_*::*sfgfp*]. STM was grown in LB broth and samples were collected in hourly intervals for quantification of DsRed fluorescence levels by flow cytometry (FC) (**Fig. S 1AB**). Without further induction, DsRed intensity decreased over time proportional to bacterial replication, validating the principle of fluorescence dilution upon bacterial replication.

For infection of the murine macrophage-like cell line RAW264.7, STM WT harboring the reporter [P*_tetA_*::*dsred* P*_msrA_*::*sfgfp*] was grown overnight (o/n) in presence of AHT to induce expression of DsRed. RAW264.7 cells were infected by DsRed-positive STM WT. At 8 h p.i. in culture without AHT, the population was released, subjected to FC and the x-median RFI of DsRed was determined (**Fig. S 1C**). Two different subpopulations were detected when using STM WT [P*_tetA_*::*dsred* P*_msrA_*::*sfgfp*] (blue histogram). One subpopulation showed normal, non-diluted DsRed intensities comparable to constitutive DsRed expression via the EM7 promoter (**Fig. S 1C**, black histogram). The second subpopulation showed a lower DsRed intensity due to FP dilution. Without addition of AHT to o/n cultures, no DsRed-positive bacteria were detected (**Fig. S 1C**, red histogram). Furthermore, NR subpopulations were detected 16 h, 24 h and 48 h p.i. using STM WT (**Fig. S 1D**), or various STM mutant strains (data not shown). We also determined the detection accuracy of the cytometer used (**Fig. S 1E-H**). The measurements demonstrated that up to 10,000-fold differences in numbers of STM expressing DsRed or sfGFP were determined very precisely.

FC analysis of the distribution of DsRed and sfGFP fluorescence of the intracellular population revealed a very small DsRed-positive NR subpopulation (app. 0.4%), compared to the DsRed-negative R subpopulation (**Fig. S 2A**). Calculating the proportion of NR STM of various mutant strains demonstrated that Δ*ssaV* and Δ*dksA* showed a higher proportion of NR STM of about 7% and 20%, respectively (**Fig. S 2B**), as result of attenuated intracellular replication of R STM. Taken together, analyses of FP dilution allowed discrimination of R and NR STM, and detection of NR STM of mutant strains at various time points p.i.

### Stress response of non-replicating intracellular Salmonella enterica

Next, we analyzed the stress response in NR compared to R STM. For the detection of the entire intracellular population, STM WT harboring [P_EM7_::*dsred* P*_xxx_*::*sfgfp*] for constitutive DsRed expression was used. For detection of intracellular NR STM, STM WT harboring [P*_tetA_*::*dsred* P*_xxx_*::*sfgfp*] for AHT-induced DsRed expression was used. STM WT harboring the various reporters were used to infect RAW264.7 macrophages (**Fig. 2**). For STM WT harboring the reporters for detection of NR bacteria, AHT was added to o/n cultures of inoculum. The bacteria were released from host cells 8 h p.i. and subjected to FC (**Fig. 2AB**), or immuno-stained against O-antigen and imaged by fluorescence microscopy (**Fig. 2C-H**). STM WT harboring a plasmid for constitutive expression of DsRed, but lacking expression of sfGFP served as negative control for adjustment of FC gating as described before (Schulte et al., 2020).

**Fig. 2:**
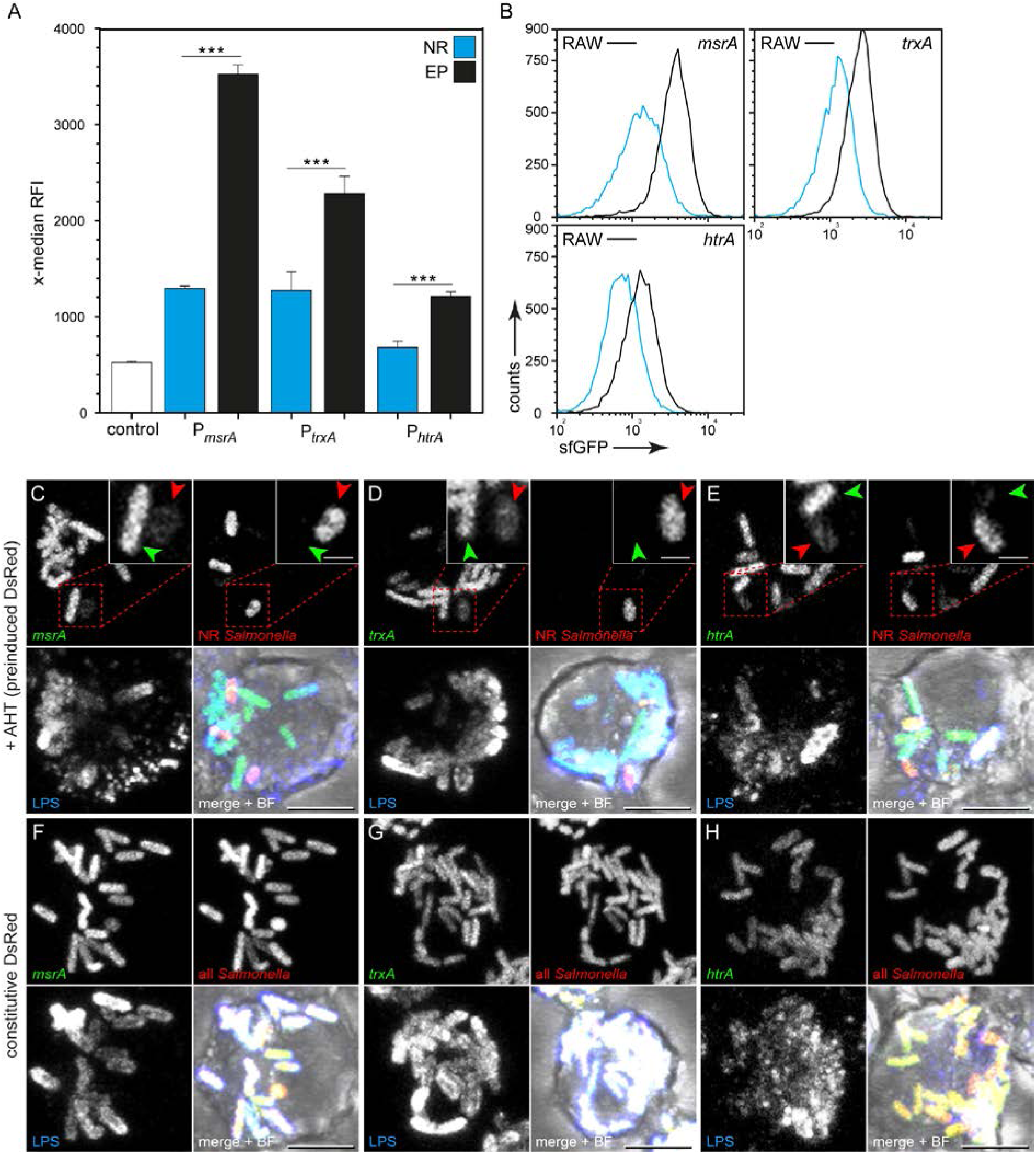
Lower levels of stress response of non-replicating STM WT compared to replicating intracellular STM. STM WT harboring AHT-inducible (NR, blue bars) or constitutive (EP, black bars) dual fluorescence reporter for *msrA, trxA* and *htrA* were grown o/n in LB medium containing 50 ng × ml^-1^ AHT. Before infection of RAW264.7 macrophages, AHT was removed. At 8 h p.i., infected cells were lysed, or fixed for FC analyses (A, B), or microscopy (C-H), respectively. For FC, released STM were recovered, fixed, and analyzed. As non-induced control, dual FP reporter plasmids were used with a frame shift mutation in *sfgfp* (described before in (Noster et al., 2019). A) The x-median represents the stress-induced sfGFP signal of the DsRed-positive intracellular bacterial population 8 h p.i. Means and standard deviations of one representative experiment are shown. B) Representative histograms corresponding to A) are shown. Statistical analysis was accomplished by SigmaPlot by One Way ANOVA and significance levels are indicated as follows: *, p < 0.05; **, p < 0.01; ***, p < 0.001; n.s., not significant. C-H) The entire population of intracellular STM was detected by immuno-staining of O antigen (blue). F-H) In addition, constitutively *dsred*-expressing STM show the entire intracellular bacterial population. C-E) NR and R STM were positive or negative for DsRed fluorescence (red), respectively. sfGFP signals (green) indicate induction of *msrA, trxA* or *htrA*. Representative NR STM and R STM are indicated by red and green arrowheads, respectively. Scale bars, 5 and 1 µm in overview and detail, respectively.

For all stress reporters investigated, i.e P*_msrA_*::*sfgfp,* P*_trxA_*::*sfgfp,* P*_htrA_*::*sfgfp*, we observed increased sfGFP levels in NR STM (**Fig. 2AB**). In contrast, the overall sfGFP expression of the entire bacterial population was higher (2.72-, 1.79-, and 1.77-fold induction of P*_msrA_*::*sfgfp*, P*_trxA_*::*sfGFP,* and P*_htrA_*::*sfGFP*, respectively, compared to NR STM). Fluorescence microscopy showed comparable results (**Fig. 2C-H**), and R and NR STM were readily distinguished by the absence or presence of DsRed fluorescence, respectively (**Fig. 2C-E**). Correlation to sfGFP fluorescence signals showed lower induction of *msrA, trxA,* or *htrA* in NR. STM WT harboring the stress reporters with constitutive DsRed expression all displayed DsRed and sfGFP fluorescence, indicating induction of *msrA, trxA* or *htrA* (**Fig. 2F-H**). The inoculum without AHT addition lacked DsRed synthesis, and accordingly DsRed fluorescence was not detectable (**Fig. S 3**). NR STM WT resided in LAMP1-positive compartments throughout intracellular presence (**Fig. S 4**), in line with prior findings (Helaine et al., 2010) and proving precision of our dual fluorescence reporter system.

We have reported that specific virulence factors of STM contribute to reduce exposure to host defense mechanisms and stressors (Noster et al., 2019; Schulte et al., 2020). Therefore, we compared the stress response of R STM to NR STM strains deficient in SPI2-T3SS or SRS in RAW264.7 macrophages at various time points p.i. (**Fig. 3**). We used P*_msrA_* as representative for induction of SRS. For all mutant strains analyzed, we observed a lower *msrA* induction of the NR subpopulation compared to the entire population at 24 h p.i. (**Fig. 3A**). We further calculated the time-resolved stress response of R and NR STM (**Fig. 3B**). The depicted slopes represent the sfGFP intensity increase over the time of infection (8 – 24 h p.i.). Higher stress induction over a constant period of time resulted in a higher slope as shown for example for the entire population of STM Δ*ssaV* and Δ*dksA* strains (red and yellow line in **Fig. 3B**). We observed that stress induction in NR STM only increases very slightly during the course of infection (dashed lines). In contrast, R STM showed a high increase of the sfGFP signal. We can conclude that intracellular NR STM perceive external stress conditions and did not encounter the same stress levels as R STM.

**Fig. 3:**
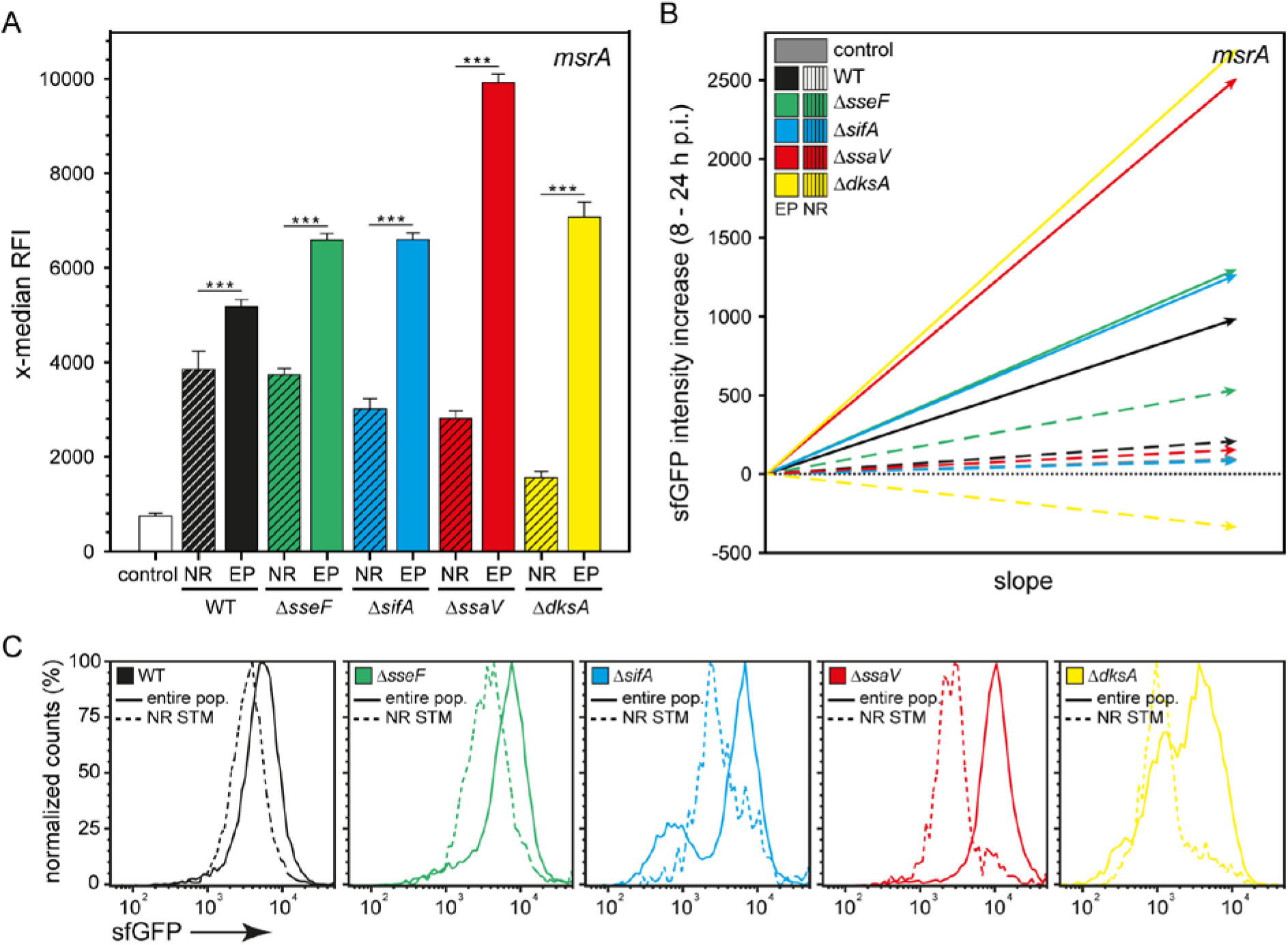
Non-replicating SPI2-T3SS or SR mutant strains also show lower stress response compared to respective subpopulation. STM WT, Δ*sseF*, Δ*sifA*, Δ*ssaV,* and Δ*dksA* strains harboring AHT-inducible (NR), or constitutive (entire population, EP) dual fluorescence reporter for *msrA* were grown o/n in LB medium. AHT was present for AHT-inducible reporters and removed before infection. RAW264.7 macrophages were infected, lysed 24 h p.i., liberated STM were recovered, fixed, and subjected to FC analyses. As negative control, a dual FP reporter plasmid with *sfgfp* inactivated by a frame-shift was used. A) The x-median represents the *msrA*-induced sfGFP signal of the DsRed-positive intracellular bacterial population at 24 h p.i. Means and standard deviations of one representative experiment are shown. B) The depicted slopes represent the sfGFP intensity increase over the time of infection. C) Representative histograms corresponding to A) are shown. Statistical analysis was performed as described for Fig. 2.

### Metabolic activity analyses of intracellular persisters at the early phase of infection

The lower induction of stress reporters in NR STM can be explained by i) lower levels of stressor acting on this population compared to R STM, or ii) reduced biosynthetic activity resulting in lower synthesis of sfGFP. To distinguish, we analyzed the metabolic activity of NR STM WT. For this, dual fluorescence reporter plasmid harboring arabinose-inducible *dsred* expression and AHT-inducible *sfgfp* expression was generated (**Fig. S 5A**). Presence of arabinose during culture of inoculum resulting in synthesis of DsRed. After removal of arabinose and infection of RAW264.7 macrophages, R STM dilute DsRed while NR STM maintain DsRed levels. sfGFP expression was induced by addition of AHT to infected cells, serving as proxy for the biosynthetic capacity (metabolic activity) of STM (**Fig. S 5BC**). For functional control, STM WT harboring the double-inducible dual fluorescence reporter [P_BAD_*::dsred* P*_tetA_::sfgfp*] was grown in LB medium and samples were collected for quantification of DsRed and sfGFP levels by FC (**Fig. S 5D-F**). Arabinose was added at the beginning of subculture (-4 h). After 4 h, the x-median RFI of DsRed increased, however, the sfGFP intensity remained constantly low. After removal of arabinose and start of a further subculture without the presence of arabinose but in the presence of AHT (0 h), the intensity of DsRed decreased within 3 h of subculture due to fluorescence dilution. In contrast, the x-median RFI of sfGFP increased (+3 h), confirming induction by AHT.

Next, we analyzed the metabolic activity of NR STM WT [P_BAD_*::dsred* P*_tetA_::sfgfp*] in RAW264.7 macrophages at 24 h p.i. (**Fig. 4**) following the experimental design depicted in **Fig. S 5C**. At 22 h p.i., AHT was added directly to the cell culture medium to induce expression of sfGFP. At 24 h p.i., bacteria were released from host cells and subjected to FC as described above. Plotting the population against their DsRed and sfGFP intensities allowed clear discrimination of the various subpopulations (**Fig. 4A**). STM positive only for DsRed represent metabolically inactive NR persisters. STM positive both for DsRed and sfGFP represent metabolically active NR persisters. STM only sfGFP-positive represent a metabolically active R population. Particles both negative for DsRed and sfGFP represent the background signal containing host cell debris. Without the addition of arabinose to o/n cultures, and without addition of AHT to infected cells, neither DsRed-nor sfGFP-positive STM could be detected by FC (**Fig. 4A**). Comparing the amount of metabolically active to inactive NR STM revealed that app. 30-40% of all persisters showed metabolic activity at 24 h p.i. (**Fig. 4B**). Comparing the metabolic activity of active persisters to active R STM showed that the metabolic capacity did not differ significantly (**Fig. 4C**), demonstrating that metabolically active NR STM showed the same metabolic activity compared to R STM.

**Fig. 4:**
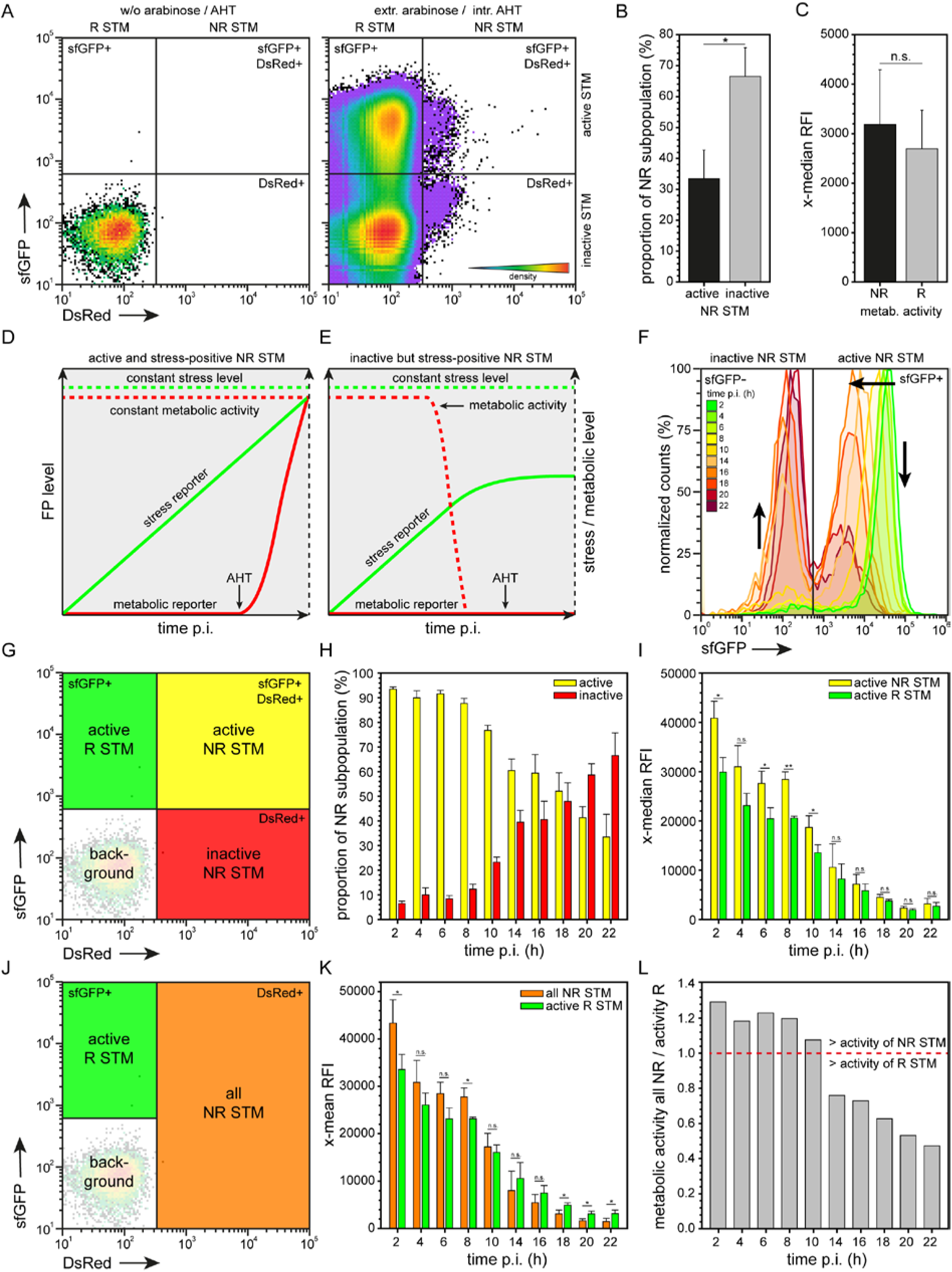
Non-replicating persisters show equal metabolic activity compared to replicating STM WT in the early phase of infection. STM WT harboring double-inducible dual fluorescence reporter was grown o/n in LB medium in the presence of arabinose. Before infection, arabinose was removed. RAW264.7 macrophages were infected, AHT was added 22 h p.i., and cells were lysed 24 h p.i. Liberated STM were recovered, fixed and subjected to FC. A) Plotting the entire intracellular bacterial population against their arabinose-induced DsRed and AHT-induced sfGFP intensity shows distinct populations. As control, STM without addition of arabinose or AHT was analyzed. B) The proportion of metabolically active and inactive NR STM WT at 22 h p.i. is shown. C) Comparison of the metabolic activity of active NR and R STM WT 22 h p.i. D-H) STM WT was cultured, infected and prepared for FC as described above. AHT was added at various time points p.i. and cells were lysed 24 h p.i. D) FP levels anticipated for metabolically active and stress signal-positive NR STM. E) Detected metabolically inactive NR STM may be stress signal-positive because of metabolic activity before addition of AHT. F) Representative histograms of the metabolic activity of intracellular NR STM WT at various time points p.i. Arrows indicate increase in metabolically inactive NR STM, and decrease in metabolically active NR STM over time of infection. G) Yellow, green, and red gates define subpopulations compared in (H) and (I). H) The proportion of metabolically active and inactive NR STM at the various time points was calculated. I) Comparison of the metabolic activity of active NR and R STM WT at various time points p.i. J) Green and orange gates define subpopulations compared in (K). K) Comparison of the metabolic activity of all NR and R STM WT at various time points p.i. L) The ratio of metabolic activity of all NR STM to metabolic activity of R STM is shown. Means and standard deviations of one representative experiment are shown. Statistical analysis was performed as described for Fig. 2.

However, if about half of the persistent population was inactive at 24 h p.i., why did we only detect a homogeneous stress-induced population instead of two subpopulations (**Fig. 2B**)? Is there one subpopulation that showed stress responses and another lacking response? To measure the metabolic activity at 24 h p.i., AHT was added 22 h p.i. If a persister is metabolically active at 22 h p.i., this cell starts sfGFP synthesis (**Fig. 4D**), but inactive persisters do not. However, a persister metabolically inactive at 22 h p.i. may have been active in the prior period of infection (**Fig. 4E**). If this holds true, the proportion of active persisters should be high at beginning of infection and decrease over time of infection, similar to result by Helaine et al. (2014) for time points 24-72 h p.i. Since stress induction is measured by accumulation of sfGFP, an active persister harboring the dual fluorescence stress reporter could synthesize sfGFP at the early phase of infection, become inactive e.g. at 10 h p.i., thus appears metabolically inactive at 24 h p.i. However, such cell would still contain sfGFP synthesized in response to stressors and should be detected as stress signal-positive at 24 h p.i. (**Fig. 4E**).

To test this hypothesis, we calculated the proportion of metabolically active and inactive persisters in the early phase of infection (**Fig. 4GH**, FC histogram shown in **Fig. 4F**). For that, AHT was added at indicated time points, however, host cells were lysed 24 h p.i. for clear separation of R and NR STM. At 2 h p.i., appr. 95% of all NR STM were metabolically active. Starting 8 h p.i., the amount of active persisters decreased constantly. The metabolic capacity of this active persistent population was always on the same level compared to active R STM (**Fig. 4I**). However, in the early phase of infection from 2-10 h p.i., the metabolic capacity of active NR STM was even higher compared to active R STM. The higher x-median RFI of sfGFP of both, active R and NR STM at the early time points post infection was resulting because all samples were lysed 24 h p.i.

We also calculated the metabolic capacity of the entire NR STM population, because when measuring stress induction of NR STM we are not able to discriminate between metabolically active and inactive NR STM (**Fig. 4JK**). In addition, we calculated the ratio of the metabolic activity of all NR compared to active R STM (**Fig. 4L**). Values above a ratio of 1 (red line) indicate higher metabolic activity of the entire NR population. The metabolic activity of the entire NR population up to 10 h p.i. was approximately at the same level as that of R STM. To conclude, because of the same activity of R and NR STM, the lower stress response of persisters 8 h p.i. is not due to a lower metabolic capacity of persisters.

### Antibiotic exposure of persisters results in decreased stress response

Since intracellular persisters are able to withstand bactericidal antibiotic exposure, we exposed STM in RAW264.7 macrophages to cefotaxime and measured stress induction of NR STM WT at 24 h p.i. As representative indicator for stress response induction P*_msrA_* was used. At 10 h p.i., 200 µg × ml^-1^ cefotaxime was added, following incubation for further 14 h. After cefotaxime treatment for 14 h, the subpopulation of R STM was highly reduced (**Fig. S 6A**). The proportion of NR STM WT increased from 0.32% to 13% (**Fig. S 6B**) representing a 41.04-fold increase (**Fig. S 6C**). However, the replicating population has not been fully removed, in contrast to prior observations (Helaine et al., 2014). Fluorescence microscopy of infected RAW264.7 macrophages at 24 h p.i. (**Fig. S 6DE**) showed signal patterns supporting the FC data. Without antibiotic treatment, STM WT replicated efficiently in macrophages also containing NR STM showing DsRed fluorescence (**Fig. S 6D**).

After cefotaxime treatment, RAW264.7 macrophages contained both, R and NR STM WT (Fig. S 6E, middle panel, indicated by red and green arrows), or NR STM WT only (**Fig. S 6E**, lower panel). Analysis of stress response showed that after antibiotic treatment the x-median RFI of sfGFP decreases, however, cefotaxime-treated NR STM WT still showed an active stress response (**Fig. 5A**). When we checked the entire intracellular population, we observed that after cefotaxime treatment the *msrA* induction also dropped to a level comparable to non-treated NR STM.

**Fig. 5:**
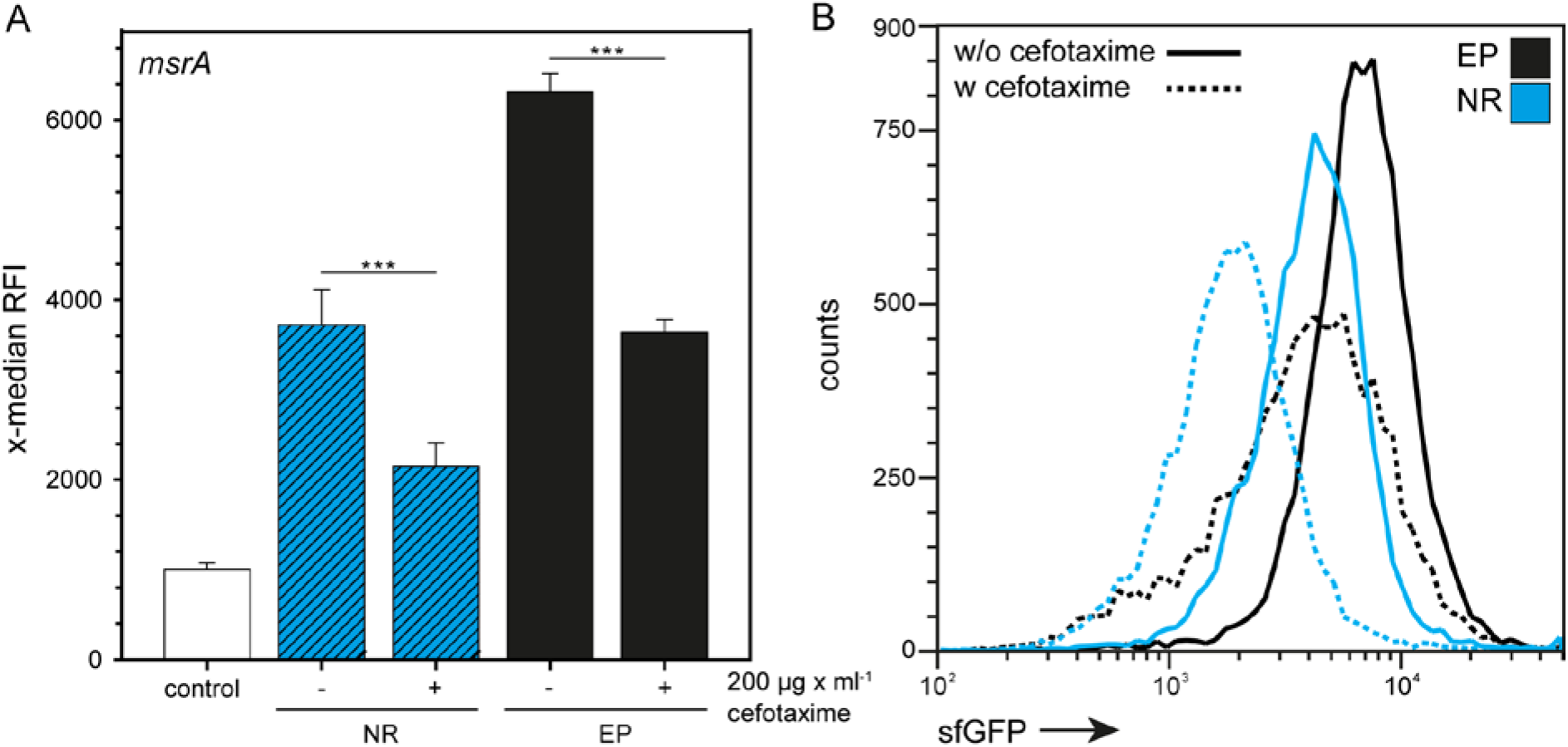
Cefotaxime-treated non-replicating STM WT show slight induction of stress response. STM WT harboring AHT-inducible (NR) or constitutive (EP) dual fluorescence reporter for *msrA* was grown o/n in LB medium, in the presence of AHT or the NR reporter. Before infection, AHT was removed. RAW264.7 macrophages were infected, cefotaxime was added to the cells at 10 h p.i. if indicated, and cell were lysed at 24 h p.i. Liberated STM were recovered, fixed and subjected to FC. As negative control, a dual FP reporter plasmid with *sfgfp* inactivated by a frame-shift was used. The x-median (A) represents the P*_msrA_*-induced sfGFP signal of the DsRed-positive intracellular bacterial population, and (B) shows a representative histogram. Means and standard deviations of one representative experiment are shown. Statistical analysis was performed as described for Fig. 2.

Since persisters are a subpopulation of NR STM that are able to resume growth after release from host cells, we controlled that we indeed analyzed growth-competent persister cells. For that, we inoculated the released population after lysis of macrophages into fresh LB medium and calculated the relative amount of DsRed-positive persisters 0, 2, 4 and 6 h after reinoculation (**Fig. S 7A**). The number of DsRed-positive events (NR persisters) did not decrease within the first 4 h after reinoculation (**Fig. S 7BC**). After that, the number of DsRed-positive events decreased to about 50%, indicating that NR persisters are still viable and competent to regrow. As determined in **Fig. S 1E-H**, ratios of up to 1:10,000 between STM DsRed and STM sfGFP were precisely determined. In addition, detection of the optical density of the reinoculated cultures showed that cefotaxime-treated STM WT started to replicate after 4-6 h (**Fig. S 7D**). Therefore, the persisters analyzed were able to regrow 6 h after reinoculation that was in line with optical density of cultures.

### Stress response of persisters within primary human phagocytes

Finally, we determined the stress response of NR STM within primary human macrophages. Primary cells were isolated from buffy coat and showed M1 polarization after differentiation (data not shown). P*_msrA_* was used as representative for stress response induction. Infection and FC analysis was performed as described above and NR STM were detected inside human macrophages at 8 h p.i. (**Fig. 6A**). Comparison of *msrA* induction of NR STM in murine and human macrophages revealed an equal x-median RFI of sfGFP (**Fig. 6B**). In addition, the *msrA* induction of the NR subpopulation [P*_tetA_*::*dsred* P*_msrA_*::*sfgfp*] and the entire population [P_EM7_::*dsred* P*_msrA_*::*sfgfp*] showed the same level in human macrophages. Plotting the bacterial population against their AHT-induced DsRed and *msrA*-induced sfGFP intensity showed that in primary human macrophages only NR STM were present, while a subpopulation of R STM was absent (**Fig. 6C**). To conclude, also in primary human macrophages NR STM WT showed an active stress response that was comparable to RAW264.7 macrophages.

**Fig. 6:**
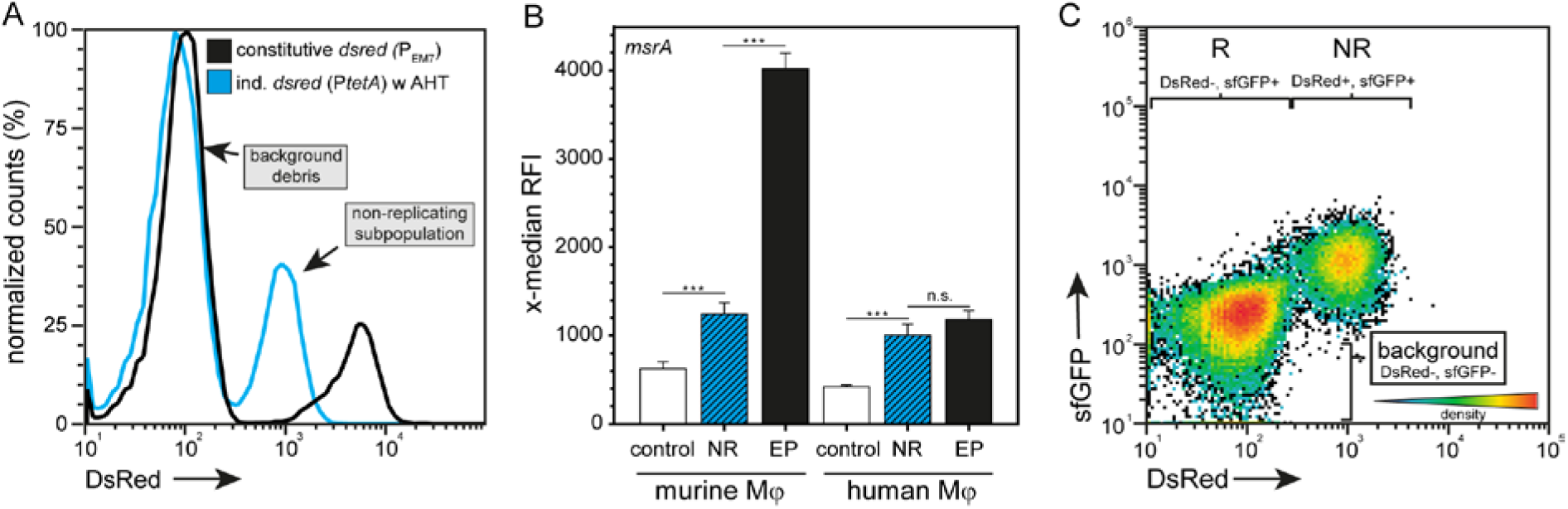
Stress response of non-replicating STM WT in primary human macrophages isolated from peripheral blood. STM WT harboring AHT-inducible (NR) or constitutive (EP) dual fluorescence reporter for *msrA* was grown o/n in LB medium in the presence of AHT if necessary. Before infection, AHT was removed. Murine RAW264.7 or human primary macrophages from peripheral blood were infected and lysed 24 h p.i. Liberated STM were recovered, fixed, and subjected to FC. As negative control, a dual FP reporter plasmid with *sfgfp* inactivated by a frame-shift was used. A) Detection of intracellular NR STM in human macrophages. B) Comparison of *msrA* induction within murine and human macrophages. The x-median represents the *msrA*-induced sfGFP signal of DsRed-positive intracellular bacteria. Means and standard deviations of one representative experiment are shown. C) STM WT only shows a NR subpopulation when plotting the bacterial population against their AHT-induced DsRed and *msrA*-induced sfGFP intensity. Statistical analysis was performed as described for Fig. 2.

## Discussion

### NR intracellular STM are capable to respond to host-imposed stressors

In this study, we further developed and applied FP-based reporters for analysis of stress response of intracellular NR STM at single cell level. This approach enables quantification of the stress response of distinct intracellular subpopulations of STM. Our study demonstrates that NR STM persister cells perceive their intracellular environment and respond to stressors by activating a stress response even under antibiotic pressure (**Fig. 7**). These cells regrew in rich medium after infection, proving that proper NR STM persister cells were examined.

**Fig. 7:**
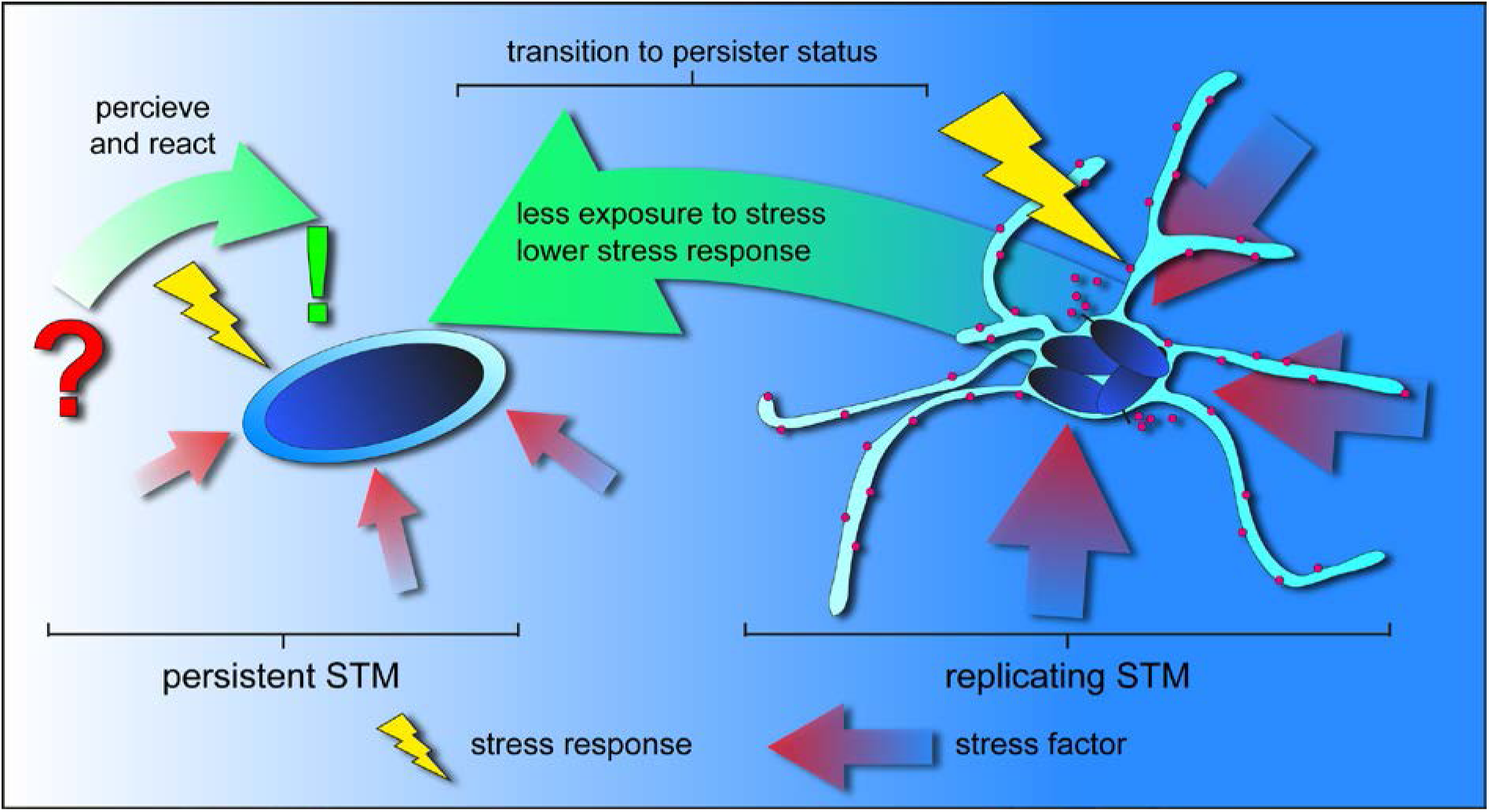
Transition to persister status results in decreases exposure to stressors and enables survival of harsh conditions in host cells. Replicating intracellular STM show a diverse stress response in reaction to harsh environmental conditions. Intracellular persistent STM are able to perceive and actively respond to stressors by turning on SRS. The induced stress response of persister is lower compared to the replicating subpopulation of intracellular STM suggesting that persister encounter lower exposure to stressors. By that, transition to persistent state inside host cells promotes stress tolerance, survival, and dissemination of the pathogen due to re-growth after subsidence of stressful conditions.

Our findings support previously reported metabolic activity of intracellular STM persisters, and translocation by T3SS-SPI2 (Helaine et al., 2014; Helaine et al., 2010; Stapels et al., 2018). STM persisters express SPI2 genes and for this, bacteria must perceive their environment to activate gene expression (Stapels et al., 2018). Our results corroborate that STM persister cells are not segregated from the intracellular habitat, but rather sense their environment and changes within. Persisters that maintain effector delivery and are able to respond to stress factors but cease to grow, still provide evolutionary benefit. Since the persister status is reversible and a single cell can give rise to a new susceptible population, the intracellular heterogeneity of *Salmonella* provides adaptive advantage and favors survival of the entire population during infection of host cells or in general, under adverse environmental conditions. Since we observed lower stress response of persister cells compared to R STM, independent from the duration of infection or function of virulence factors, we conclude that the persistent status serves as a kind of niche by protecting NR STM from severe stress exposure (**Fig. 7**). The host cell may no longer be able to exert stress on NR STM persister cells, which in turn results in lower stress response. Beyond this, STM persisters reside individually within segregated SCV during the whole course of infection, further promoting their protective niche. This compartment shields STM from the cytosolic environment and host cell defense mechanisms such as xenophagy (reviewed in Gomes and Dikic, 2014). In addition, the SCV of NR STM is separated from the SCV/SIF continuum containing R STM (Liss et al., 2017).

### At initial infection, a high proportion of NR STM is metabolically active

We observed that at initial infection of macrophages, the proportion of metabolically active NR STM was as high as 95%. Thus, the lower stress response of NR STM is not a consequence of a lower metabolic capacity. Starting about 8 h p.i., the proportion of active persisters constantly decreased. This is in line with previous observations that detected about 70% of active NR STM in RAW264.7 macrophages 12 h p.i., and about 25% active NR STM at 24 h p.i. (Helaine et al., 2010). Our studies extended these analyses to early time points p.i. In addition, we compared the level of metabolic activity, which was comparable to R STM until 10 h p.i. if we average the total activity of the entire NR population. If only metabolically active NR STM persisters are considered, these did not show a significantly different activity compared to active R STM over the entire period of infection. If a small reservoir of persisters always remains active over a longer period of time, this may allow formation of new populations in the host, and thus to trigger relapses. The metabolic activity of NR STM at the beginning of infection may serve to successfully establish their persistent status. For *M. tuberculosis* or *E. coli*, *in vitro* transcriptome studies indicated that persisters showed downregulation of metabolic genes and therefore decreased metabolism (Keren et al., 2011; Shah et al., 2006). Genes with functions in energy production and TCA cycle were downregulated in *Burkholderia cenocepacia* persisters (Van Acker et al., 2013). For intracellular STM, however, there is evidence that distinct subpopulations of metabolically active and inactive persisters exist (Helaine et al., 2010; Stapels et al., 2018). The observation of metabolically active NR bacteria is not limited to *Salmonella* species, but also reported for *M. tuberculosis* in macrophages, and could therefore be a widespread phenomenon (Manina et al., 2015).

When examining persistent bacteria, it is highly important which time point is analyzed, and to be aware that overall metabolic activity of the NR population changes over time. This refers to the decreasing proportion of metabolically active persisters over time, and the associated activity of the entire population on average. In many studies, late time points were selected to investigate persisters, at which a small proportion of metabolically active persisters was detected, which further decreased (Helaine et al., 2010). Our studies are in line with these results and our data also showed that when the entire NR subpopulation is considered, a decreased metabolic activity from 14 h p.i. was observed. This is also in line with a model of reduced metabolism in persisters cells. Importantly, before 14 h p.i. the entire population of NR STM persisters on average does not show a reduced metabolism and the metabolic activity is not lower at any time point analyzed if only metabolically active NR STM are considered. However, since the proportion of metabolically active persisters decreases over time, the metabolic activity of the entire population of NR STM on average decreases accordingly. In turn, a reduced metabolic activity is measured when the entire population of NR STM is observed. Conventional transcriptome analyses or other population-wide approaches average the entire NR population and fail to distinguish between active and inactive persisters. Single cell analyses that are capable of addressing small subpopulations of intracellular STM, such as the approaches applied here, provide a more precise insight into the physiological state of persisters. Novel single cell transcriptomics approaches for bacteria are emerging (Imdahl et al., 2020) and may provide a future option for analyses of persisters.

### Stress response of NR STM in non-permissive primary human macrophages

We investigated stress response of NR STM persisters in much less permissive human phagocytic cells. We observed that the level of stress response triggered human macrophages was comparable to that of STM in RAW264.7 macrophages being rather permissive for STM intracellular proliferation. These results suggest that the induced stress response of NR STM is either independent of the killing capacity of the host cell, or because persisters are so well protected that even phagocytes with high antimicrobial activity do not induce higher levels of stress response.

Typhoidal serovars (TS) of *S. enterica* cause systemic infections in humans such as typhoid fever, and persistent or recurrent forms of the disease are frequent (Gotuzzo et al., 1987). Advanced analyses of stress response shall be extended to persister cells of typhoidal serovars Typhi and Paratyphi A to gain insights into their physiological state, and to test the ability to perceive stress factors. As part of the ‘stealth strategy’ (Dougan and Baker, 2014), persister formation in typhoidal serovars may occur at higher frequency. In relation to heterogeneity, it will be of interest if intracellular Typhi and Paratyphi A show distinct subpopulations with different levels of stress response. Knowledge of persister cell frequency, the proportion of metabolically active and inactive persisters, and the level of stress response of TS in comparison to STM may lead to new insight into patho-mechanisms of TS and recurrent infections. This will require further analyses in other cell lines like human phagocytic cell line U937 or primary phagocytic cells.

### Stress response of persisters and new therapeutic options?

Since persisters that emerge during infection are extremely well protected and the clinical relevance of relapsing infection due to persister is high, new strategies to eliminate persister cells are of utmost therapeutic importance. Current approaches include prevention of persister formation, identification of antimicrobial compounds that act on persister cells, or resuscitation of persister cells to restore sensitivity to conventional antibiotics (reviewed in Defraine et al., 2018). Promising studies have been performed to identify compounds acting on bacterial persister cells, however further action is required. High-throughput screening methods are often applied, but many screens were performed using planktonic persisters *in vitro*. Recent work showed that the efficacy of drugs against tuberculosis is significantly different when applied *in vivo* compared to *in vitro* conditions (Liu et al., 2016). This may also apply to other drugs, making analyses of effects on persister cells during interplay with their host essential. One approach could be host-directed therapies focusing on host-pathogen interactions, as already has been demonstrated for *M. tuberculosis* infections by treating infections with small molecules to interfere with host responses or persistence (Kim and Yang, 2017; Kiran et al., 2016). Antibody-antibiotic conjugates are another emerging approach. Antibiotics are coupled to antibodies against a specific pathogen to increase the efficiency of antibiotics against intracellular pathogens including persisters (Mariathasan and Tan, 2017). Zhou et al. (2016) showed that *S. aureus* bacteremia in mice was successfully treated, hence, this approach could serve as a novel therapeutic platform in the future. Our finding of the continuing function of SRS in persister cells and the ability of persisters to sense stressors may lead to new options for resuscitation of persisters (examples in Song and Wood, 2020). If deregulation of SRS by decoy compounds is possible, this may lead to re-initiation of normal growth and increased antibiotic susceptibility. Approaches to interfere with quorum sensing for interbacterial communication (reviewed in Defoirdt, 2018) or biofilm matrix production (Dieltjens et al., 2020) enabled new antimicrobial strategies. In contrast, potential interference with SRS in persisters needs to target single cells in complex populations within host organisms.

### Conclusions and outlook

In summary, we have developed and applied new dual FP reporters that enable extremely sensitive analyses of stress response and metabolic activity of NR persister cells. Of particular interest is that NR STM perceive their environment and display a lower stress response with constant metabolic activity during the early phase of infection. Our findings provide deeper knowledge that besides the ability to subvert immune defenses of the host, NR STM persister cells maintain capability to sense stressors and react to stress. These persisters exhibit constant metabolic activity at the beginning of the infection, which supports the pathogen to create a more permissive environment for recrudescent infections.

## Acknowledgements

This work was supported by the DFG though grants in SFB 944, projects P4 and P15. We kindly acknowledge intramural funding by profile line P2: Integrated Science of the University Osnabrück. We like to thank the Hans-Mühlenhoff-Stiftung for support of KO. We thank Tatjana Reuter and Jennifer Röder for isolation of human macrophages.

## Materials and Methods

### Generation of reporter plasmids

For the generation of a reporter plasmid to detect non-replicating STM, plasmid p5084 (P_EM7_::*dsred* P*_msrA_*::*sfgfp*), p5085 (P_EM7_::*dsred* P*_trxA_*::*sfgfp*) and p5055 (P_EM7_::*dsred* P*_htrA_*::*sfgfp*) with a constitutive expression of Dsred and *msrA, trxA* or *htrA* regulated expression of sfGFP was used. Via Gibson Assembly (GA) of PCR fragments, the EM7 promoter was replaced by the tet-ON cassette to enable artificial induction of DsRed by anhydrotetracycline (AHT).

For the generation of a single FP reporter plasmid to detect non-replicating STM, plasmid p5204 (P_EM7_::*dsred* P*_cypD_*::*sfgfp (frameshift*)) with a constitutive expression of *dsred* and a frameshift mutation in *sfgfp* was used. Via GA, the EM7 promoter was replaced by the tet-ON cassette to enable artificial induction of *dsred* by AHT.

For the generation of a dual fluorescence vitality reporter plasmid to detect the metabolic activity of non-replicating STM, plasmid p4928 (P_EM7_::*tagrfp-T* P*_tetA_*::*sfgfp*) with a constitutive expression of *tagrfp-T* and an AHT-inducible expression of *sfgfp* was used. Via GA, the EM7 promoter was replaced by *araC* P_BAD_ cassette to enable artificial induction of *tag-rfpT* by arabinose. Subsequently, p5419 (P_BAD_::*tagrfp-T* P*_tetA_*::*sfgfp*) was used to amplify the P_BAD_ and tet-ON cassette. Via GA, the EM7 promoter and the *msrA* promoter of p5084 (P_EM7_::*dsred* P*_msrA_*:*sfgfp*) was replaced by the P_BAD_ and tet-ON cassette to enable artificial induction of *dsred* by arabinose and *sfgfp* by AHT. The resulting single and dual FP reporter plasmids are listed in Table S 1 and oligonucleotides used for construction are listed in Table S 3.

### Bacterial strains, growth conditions AHT and arabinose induction

*Salmonella enterica* sv. Typhimurium strain NCTC12023 (STM) was used as wild-type strain and isogenic mutant strains used in this study are listed in Table S 2. Bacteria were cultured in lysogeny broth (LB) at 37 °C overnight (o/n) using a roller drum at 60 rpm with aeration. For maintenance of plasmids, carbenicillin was added at 50 µg × ml^-1^ as a selection marker. For induction of expression of P*_tetA_*-controlled dual FP reporter (p5205, p5300, p5202 or p5418) or P_BAD_-controlled dual FP reporter (p5426) AHT or arabinose always was directly added to LB to a concentration of 50 ng × ml^-1^ or 13.3 mM, respectively, in the o/n culture and removed when necessary by centrifugation for 3 min at 5,000 × g and washing with fresh LB.

### Isolation of primary human macrophages

Buffy coat was obtained from pooled samples of voluntary, anonymous blood donors via the blood bank of Deutsches Rotes Kreuz (Springe, Germany). Lymphocytes were prepared from buffy coat by Ficoll-Hypaque density gradient centrifugation (see Bonifacino et al., 2004). Buffy coat was diluted in PBS (ratio 1:3) and Ficoll-Hypaque was added, following centrifugation for 20 min at 800 *x g*. Afterwards, 5 × 10^8^ cells were cultured in Roswell Park Memorial Institute (RPMI) medium containing 5.5 g × l^-1^ NaCl, 5.0 mg × l^-1^ phenol red, 2.0 g × l^-1^ NaHCO_3_, 25 mM HEPES, 4 mM stable glutamine without sodium pyruvate (Biochrom), 100 units × ml^-1^ penicillin, 100 µg × ml^-1^ streptomycin, 2.5 ng × ml^-1^ GM-CSF (granulocyte-macrophage colony-stimulating factor) and supplemented with 10% human plasma. After o/n incubation at 37 °C in an atmosphere of 5% CO_2_ and 90% humidity, non-adherent cells were removed and adherent cells were cryo-preserved in inactivated fetal calf serum (iFCS) and DMSO.

### Cell lines and cell culture

For infection experiments murine RAW264.7 macrophages (American Type Culture Collection, ATCC no. TIB-71), RAW264.7 macrophages stably transfected with LAMP1-GFP or primary human macrophages were used. RAW264.7 macrophages were cultured in Dulbecco’s modified Eagle’s medium (DMEM) containing 3.7 g × l^-1^ NaHCO_3_, 4.5 g × l^-1^ glucose, 4 mM stable glutamine without sodium pyruvate (Biochrom) and supplemented with 6% iFCS (Sigma-Aldrich). Primary human macrophages were cultured in RPMI medium containing 5.5 g × l^-1^ NaCl, 5.0 mg × l^-1^ phenol red, 2.0 g × l^-1^ NaHCO_3_, 25 mM HEPES, 4 mM stable glutamine without sodium pyruvate (Biochrom) and supplemented with 10% iFCS (Sigma-Aldrich).

### Host cell infection (gentamicin protection assay) for cytometry

Before infection, RAW264.7 or primary human macrophages were seeded in surface-treated 6- well plates (TPP) to reach confluency (∼ 2 × 10^6^ cells per well) on the day of infection. For infection, *Salmonella* strains were grown o/n in LB broth (app. 18 h). Infection was performed with a multiplicity of infection (MOI) of 10. Bacteria were centrifuged onto the cells for 5 min at 500 *x g* and infection proceeded for 25 min at 37 °C in an atmosphere of 5% CO_2_ and 90% humidity. Afterwards, infected cells were washed thrice with PBS and incubated for 1 h with cell culture medium containing 100 µg × ml^-1^ gentamicin (Applichem) to kill non-phagocytosed bacteria. Afterwards, the cell culture medium was replaced by medium containing 10 µg × ml^-1^ gentamicin until the end of the experiment. If cefotaxime was used during infection experiments, cells were washed with PBS before addition of cell culture medium containing 200 µg × ml^-1^ freshly dissolved cefotaxime.

### Host cell infection for microscopy

RAW264.7 or LAMP1-GFP RAW264.7 macrophages were seeded in surface-treated 24-well plates on glass cover slips to reach 80% confluency (ca. 3.6 × 10^5^ cells per well) on the day of infection. Cells were infected with STM strains as described above with a MOI of 50 for 8 or 24 h. Afterwards, the cells were washed thrice with PBS and fixed with 3% PFA in PBS for 15 min at RT. After that, cells were directly prepared for subsequent immunostaining.

### Immunostaining and imaging

Immunostaining of intracellular STM was performed as described before (Müller et al., 2012). After fixation of cells with 3% PFA in PBS cells were washed thrice with PBS and directly incubated in blocking solution containing 2% goat serum, 2% bovine serum albumin and 0.1% saponin in PBS for 30 min at RT. STM was stained with anti-*Salmonella* O-antigen group B factors 1, 4, 5, 12 (BD Difco, diluted 1:500 in blocking solution) for 1 h at RT. Subsequently, cells were washed thrice in drops of PBS following incubation with secondary antibody Cy5-coupled goat anti rabbit IgG (Jackson Immuno Research, 1:1,000 in blocking solution) for 1 h at RT in the dark. Afterwards, cells were washed thrice, mounted with Fluoroprep (bioMérieux) and sealed with Entellan (Merck). Fluorescence imaging was performed using the confocal laser-scanning microscope Leica SP5. Image acquisition was performed using the 100x objective (HCX PL APO CS 100×; numerical aperture: 1.4 to 0.7) and the polychroic mirror TD 488/543/633 for the three channels GFP, DsRed, and Cy5 (Leica, Wetzlar, Germany). For setting adjustment, image acquisition and image processing the software LAS AF (Leica, Wetzlar, Germany) was used.

### Flow cytometry analysis

FC of liberated STM from host cells was performed as described before (Noster et al., 2019). Briefly, FC was performed on an Attune NxT instrument (Thermo Fischer Scientific) at a flow rate of 25 μl × min^-1^. At least 10,000 bacteria were gated by virtue of the constitutive/induced DsRed fluorescence. Per gated STM cells, the intensity of the sfGFP fluorescence was determined and x-medians for the sfGFP intensities were calculated.

For the measurement of liberated bacteria (replicating and non-replicating) from host cells, infected cells were lysed at the indicated time points by 0.5% Triton X-100 in PBS for 10 min at RT with shaking to release the intracellular bacterial population. The lysate was transferred to a test tube and after pelleting of host cell debris by centrifugation for 5 min at 500 *x g*, bacteria were recovered from supernatant. Bacteria were further centrifuged for 5 min at 20,000 *x g* and fixed in 3% PFA in PBS for 15 min at RT. After fixation and a further centrifugation step, fixed bacteria were resuspended in 250 µl 100 mM NH_4_Cl in PBS for quenching of residual free aldehydes. After that, samples were directly subjected to FC.

For detection of regrowth of cefotaxime-treated intracellular non-replicating STM, the whole liberated bacterial population was reinoculated into 3 ml fresh LB medium after the bacterial supernatant was transferred to a clean test tube (Fig. S 7) and incubated at 37 °C using a roller drum at 60 rpm with aeration. At time point 0 h, 2 h, 4 h and 6 h post re-inoculation samples were taken, diluted with PBS and fixed with 3% PFA in PBS for 15 min at RT. Then, PFA was removed by centrifugation for 5 min at 20,000 *x g* and the pellet was resuspended in 250 µl of 100 mM NH_4_Cl in PBS for quenching of residual free aldehydes. After that, samples were directly subjected to FC and amounts of DsRed-positive events (non-replicating STM) was measured and depicted by events per µl.

To measure the minimal detectable threshold of bacterial events at the cytometer, various mixed ratios of constitutive DsRed and sfGFP fluorescent STM were prepared. For that, the optical density (OD_600_) of overnight cultures of STM [p5204] and STM [pWRG167] was determined following dilution of the strains to an OD_600_ of 1 (predicted amounts of bacteria: 1.1 × 10^9^ bacteria × ml^-1^). Afterwards, serial dilutions were prepared and mixed ratios of red and green fluorescent STM were prepared in PBS. Either a constant high amount of green fluorescent STM (app. 2,000 events × µl^-1^) mixed with an equal, 10-, 100-, 1,000-, or 10,000-fold reduced amount of red fluorescent STM or a constant high amount of red fluorescent STM (app. 4,000 events × µl^-1^) mixed with an equal, 10-, 100-, 1,000-, or 10,000-fold reduced amount of green fluorescent STM was measured by FC. The relative amount of green and red fluorescent STM (events × µl^-1^) measured when the equal amount of green and red fluorescent STM was present in the sample was set to 100%. As controls, only green, red or no fluorescent STM were measured.

## Suppl. Tables

**Table S1.** Plasmids used in this study

| 810 | Plasmid | Relevant genotype* | Source/reference |
| --- | --- | --- | --- |
| 811 | p5204 | P<sub>EM7</sub>::*dsred* P<sub>cypD</sub>::*sfgfp* (frameshift) | (Noster et al., 2019) |
| 812 | pWRG167 | P<sub>EM7</sub>::*sfgfp* | (Bender et al., 2013) |
| 813 | p4928 | P<sub>EM7</sub>::*tagrfp-T* P<sub>tetA</sub>::*sfgfp* | (Schulte et al., 2020) |
| 814 | p5055 | P<sub>EM7</sub>::*dsred* P<sub>htrA</sub>::*sfgfp* | (Schulte et al., 2020) |
| 815 | p5084 | P<sub>EM7</sub>::*dsred* P<sub>msrA</sub>::*sfgfp* | (Noster et al., 2019) |
| 816 | p5085 | P<sub>EM7</sub>::*dsred* P<sub>trxA</sub>::*sfgfp* | (Noster et al., 2019) |
| 817 | p5205 | P<sub>tetA</sub>::*dsred* P<sub>msrA</sub>::*sfgfp* | This study |
| 818 | p5300 | P<sub>tetA</sub>::*dsred* P<sub>htrA</sub>::*sfgfp* | This study |
| 819 | p5302 | P<sub>tetA</sub>::*dsred* P<sub>trxA</sub>::*sfgfp* | This study |
| 820 | p5426 | P<sub>BAD</sub>::*dsred* P<sub>tetA</sub>::*sfgfp* | This study |
| 821 | p5418 | P<sub>tetA</sub>::*dsred* P<sub>cypD</sub>::*sfgfp* (frameshift) | This study |
| 822 | p5419 | P<sub>BAD</sub>::*tag-rfp-T* P<sub>tetA</sub>::*sfgfp* | This study |

\* all plasmids confer resistance to carbenicillin

**Table S2.**
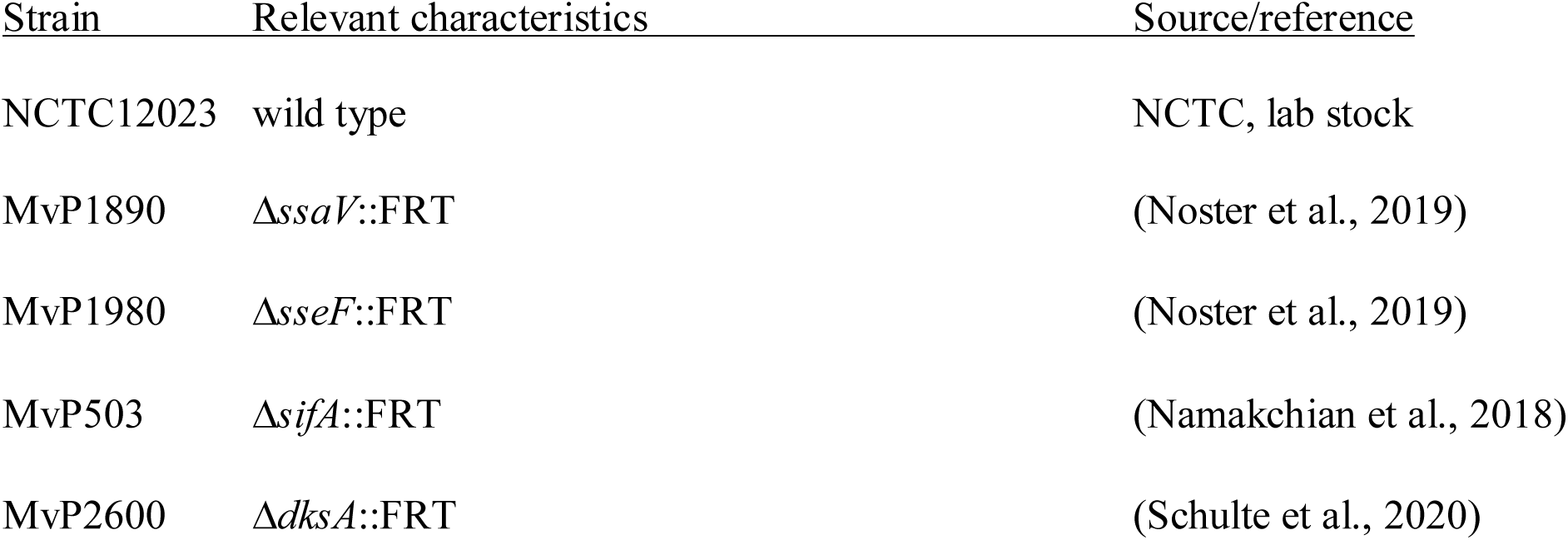
*Salmonella enterica serovar* Typhimurium strains used in this study

**Table S3.**
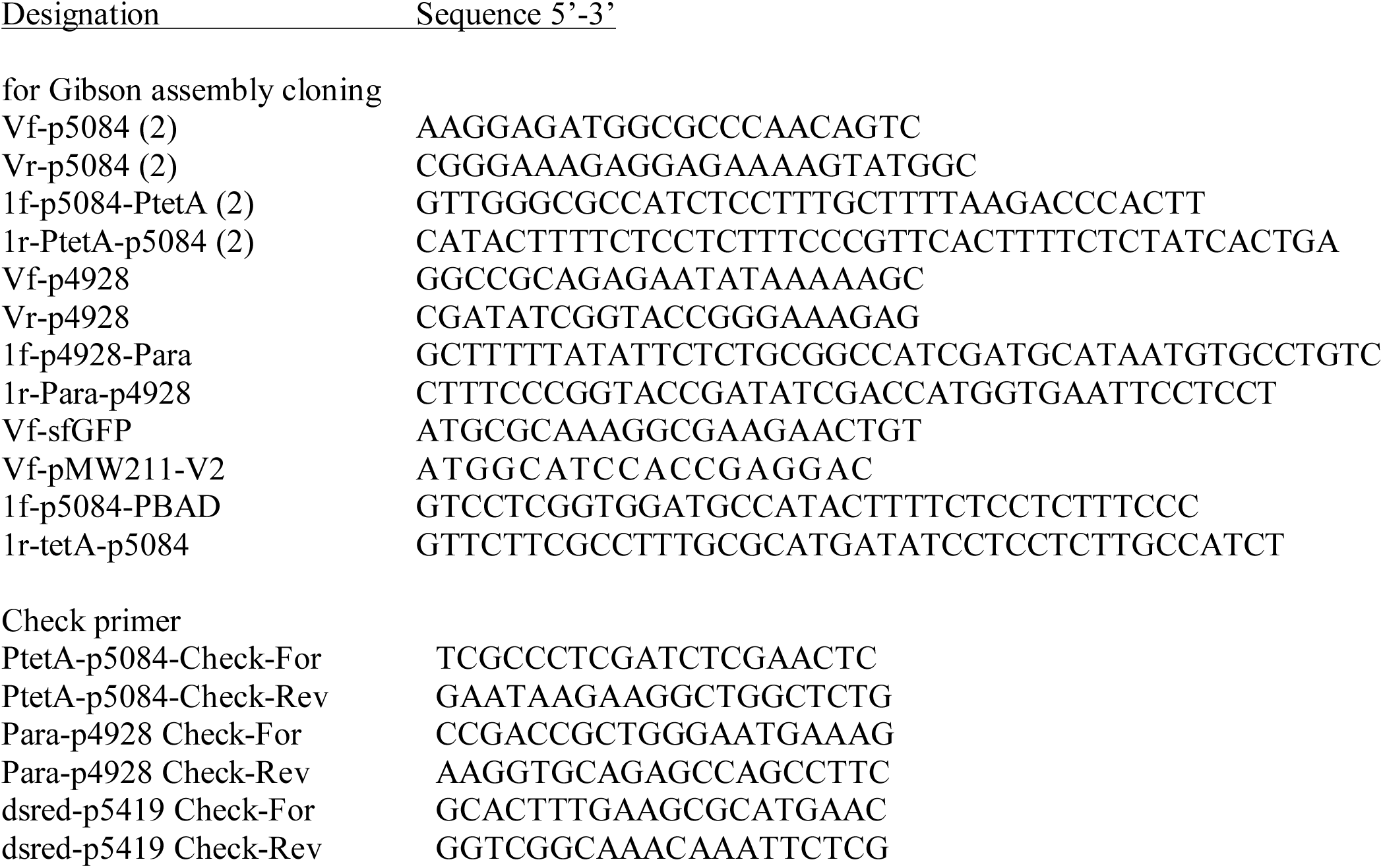
Oligonucleotides used in this study

## Suppl. Figures and Figure Legends

**Fig. S1:**
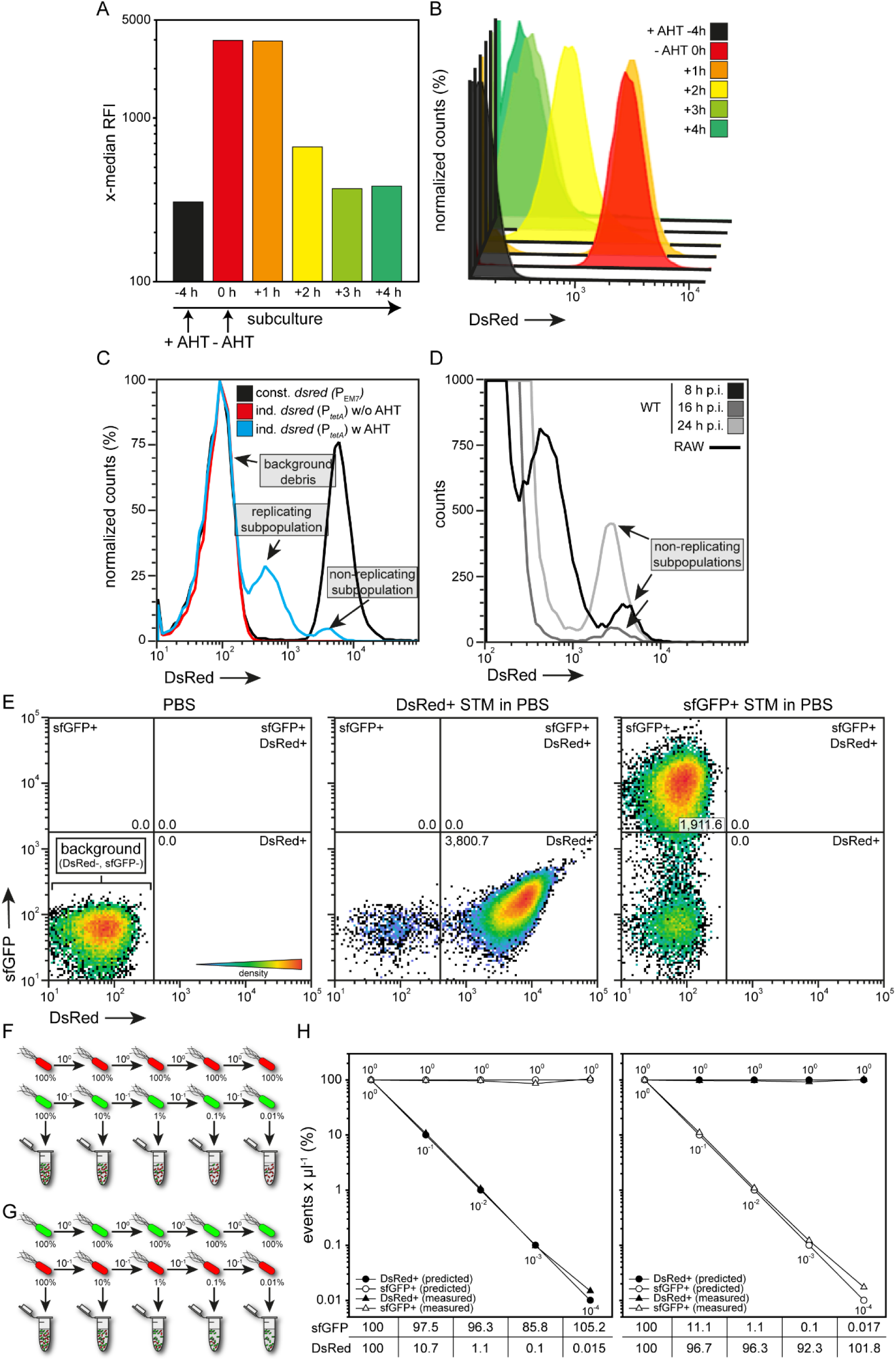
A dual fluorescence reporter suitable to detect the non-replicating subpopulation of intracellular STM. A) STM WT harboring dual fluorescence reporter with AHT-inducible *dsred* expression and *msrA*::*sfgfp* was grown in LB medium o/n and diluted in fresh LB medium containing AHT for further subculture. After 4 h of subculture, AHT was removed by centrifugation and washing. Subsequently, bacteria were subcultured in fresh LB without AHT. At indicated time points samples were taken, fixed and subjected to FC. The x-median represents the AHT-induced DsRed intensity of the entire bacterial population, and the respective histograms (B). C) STM WT harboring AHT-inducible or constitutive dual fluorescence reporter for *msrA* was grown o/n in LB medium in the presence of AHT if necessary. Before infection, AHT was removed. RAW264.7 macrophages were infected, and lysed 8 h p.i. Liberated STM were recovered, fixed and subjected to FC. As negative control, AHT was omitted from o/n cultures. The histogram shows that intracellular NR STM can be detected inside RAW264.7 macrophages showing the same DsRed intensity compared to intracellular STM WT harboring the constitutive dual fluorescence reporter for *msrA*. If AHT was omitted, no DsRed-positive STM can be detected. D) NR subpopulations of intracellular STM WT can be detected after various time points post infection. The same experiments were performed for the AHT-inducible dual fluorescence reporter for *trxA* and *htrA,* with the same outcome (data not shown). E-H) Detection accuracy of bacterial particles on an Attune NxT cytometer. STM WT strains constitutively expressing *dsred* or *sfgfp* were grown in LB medium o/n. E) Controls of PBS without STM, only DsRed-positive STM, or only sfGFP-positive STM are shown. Events per µl are indicated in each gate. Then, mixed ratios of DsRed- and sfGFP-expressing STM were prepared in PBS. Either a constant high amount of sfGFP-positive STM mixed with an equal, 10, 100, 1,000, or 10,000-fold reduced amount of DsRed-positive STM (F), or a vice versa (G) was prepared and directly subjected to FC. H) The relative amount (events × µl^-1^) of sfGFP- and DsRed-positive STM measured in the first sample was set to 100% (10^0^/10^0^). Mixtures with gradual 10-fold reduction of red or green fluorescent STM within a sample containing constant high amount of green or red fluorescent STM, respectively, were quantified. The measured value (events × µl^-1^ in %) compared to the first sample containing high amounts of both, red and green fluorescent STM, is indicated below.

**Fig. S2:**
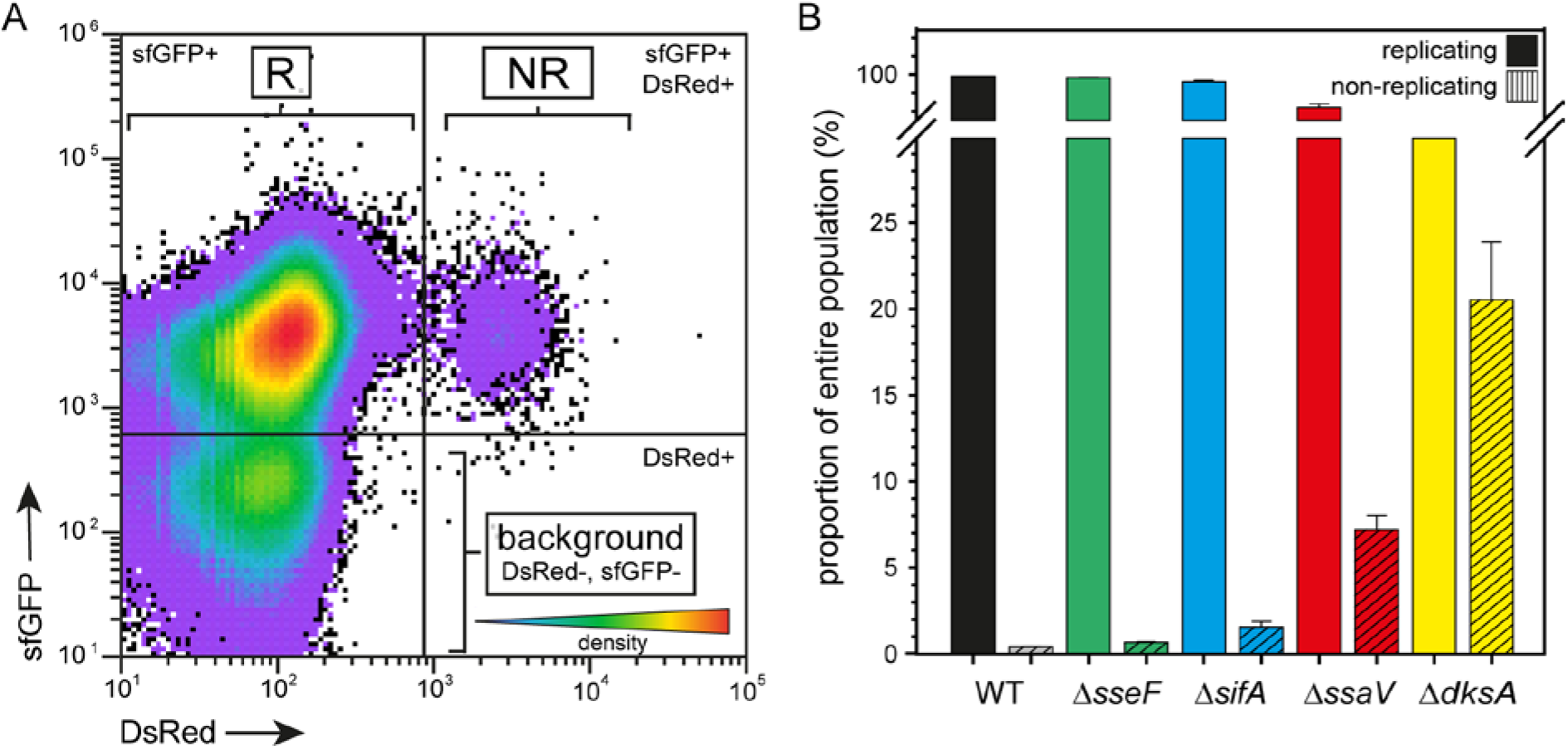
A minor fraction of the intracellular population consists of non-replicating STM. STM WT, Δ*sseF*, Δ*sifA*, Δ*ssaV* and Δ*dksA* harboring AHT-inducible dual fluorescence reporter for *msrA* were grown o/n in LB medium in the presence of AHT. AHT was removed prior to infection. RAW264.7 macrophages were infected and lysed 24 h p.i. STM released from host cells were recovered, fixed and subjected to FC. A) Analyses of the entire intracellular bacterial population for AHT-induced DsRed and *msrA*-induced sfGFP intensity shows three different populations. DsRed-negative (DsRed-) and sfGFP-negative (sfGFP-) events represent the background signal consisting of host cell debris. DsRed- and sfGFP+ events represent the replicating (R) subpopulation of intracellular STM and DsRed+ and sfGFP+ events represent the non-replicating (NR) subpopulation. B) Gating on, and quantification of R and NR subpopulations indicated the small size of the population of NR intracellular STM WT.

**Fig. S3:**
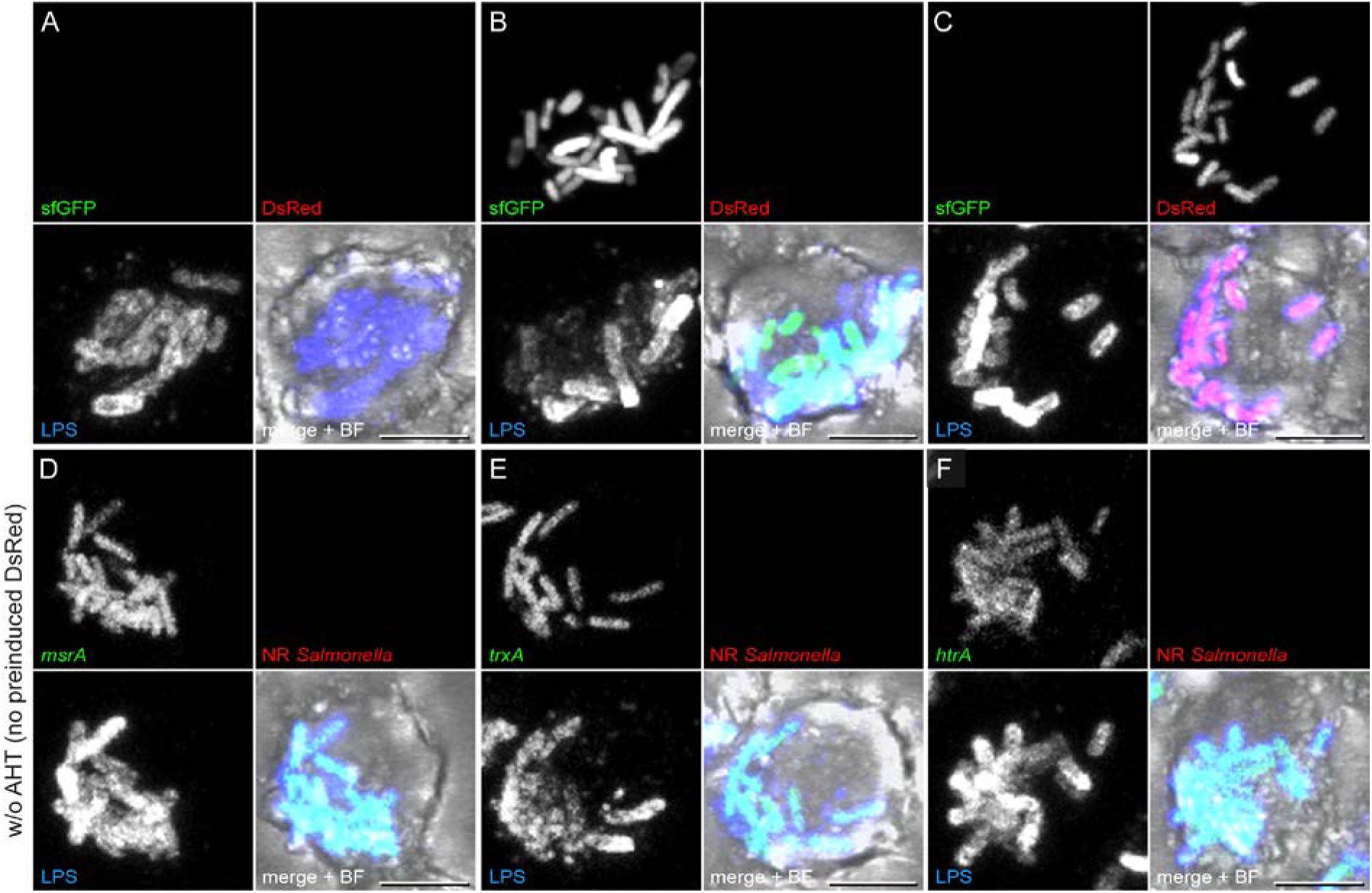
Functional control of AHT-inducible dual fluorescence reporter without AHT induction. A-F) STM WT harboring AHT-inducible (NR) dual fluorescence reporter for *msrA, trxA,* or *htrA* were grown o/n in LB medium without AHT. RAW264.7 macrophages were infected and fixed 8 h p.i. for fluorescence microscopy. STM were immuno-stained against O antigen (blue). As fluorescence controls, STM WT without expression of any FP (A), STM WT constitutively expressing *sfgfp* (B), and STM WT constitutively expressing *dsred* (C) were used. Induction of P*_msrA_* (D), P*_trxA_* (E) or P*_htrA_* (F) is shown in green. NR STM were not detected due to absence of AHT in o/n culture. Representative cells are shown. Scale bar, 5 µm.

**Fig. S4:**
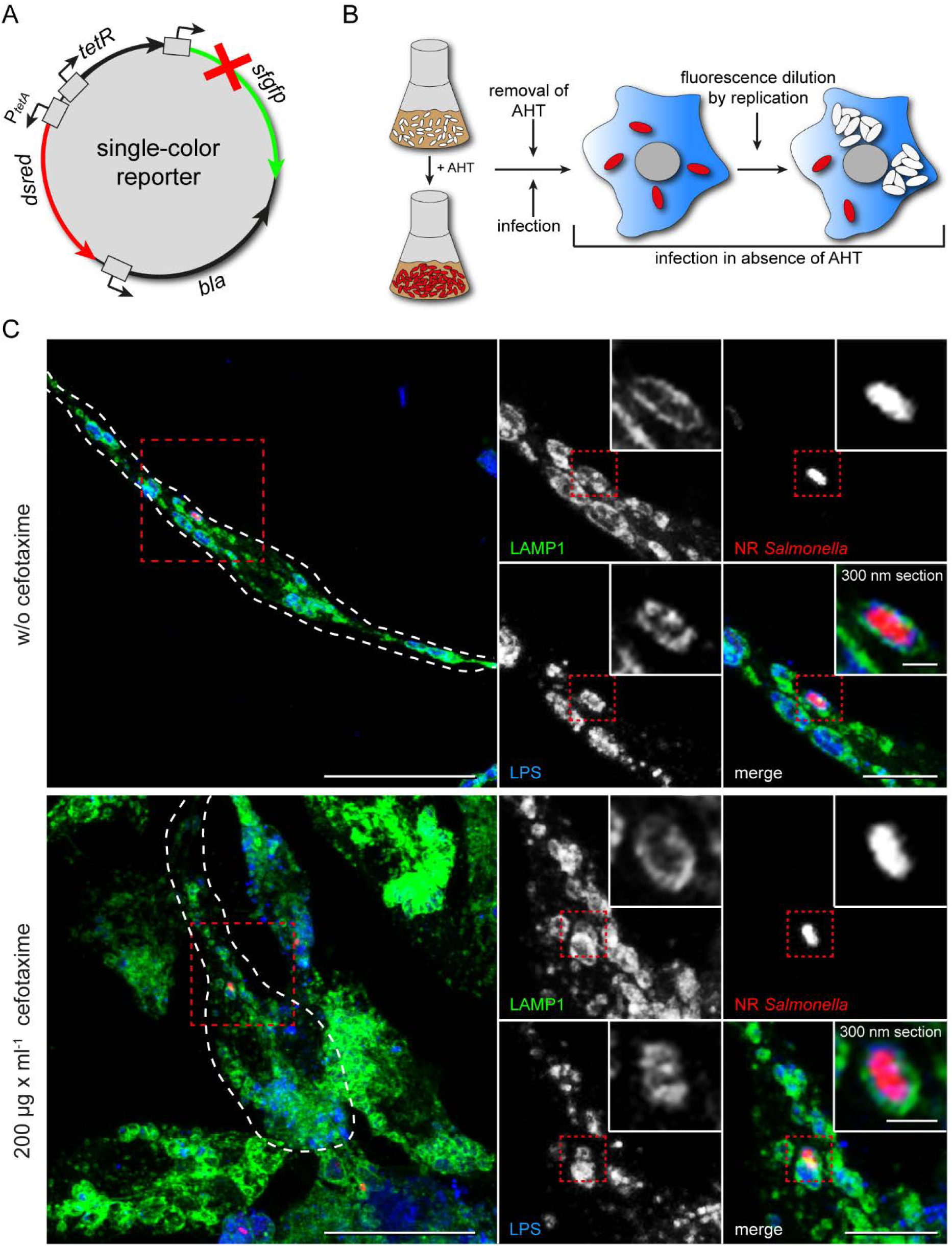
Non-replicating persisters reside inside SCV in infected host cells. A) For the analysis of NR intracellular STM, the EM7 promoter of the non-induced *sfgfp* frameshift control plasmid was replaced by the tet-ON cassette to be able to experimentally induce the expression of *dsred* by AHT. B) When adding AHT to a culture of STM growing in LB medium, bacteria start to synthesize DsRed. Before infection of macrophages, AHT is removed by centrifugation and washing. Infection is performed without addition of AHT to the cell culture medium. R intracellular STM lose DsRed via fluorescence dilution. NR intracellular STM keep the DsRed inside the cytoplasm and can be detected as DsRed-positive within e.g. RAW264.7 LAMP1-GFP macrophages during microscopy. C) STM WT harboring AHT-inducible single FP reporter was grown o/n in LB medium in the presence of arabinose. Before infection, AHT was removed. RAW264.7 LAMP1-GFP macrophages were infected, 10 h p.i. cefotaxime was added to the cells if indicated and fixed 24 h p.i. for fluorescence microscopy. Fixed intracellular STM were immuno-stained against O antigen (blue). LAMP1 is shown in green, NR STM show and R STM do not show red fluorescence. Representative cells are shown. Scale bars, 20, 5, 1 µm in overview, details, and zoom-in, respectively.

**Fig. S5:**
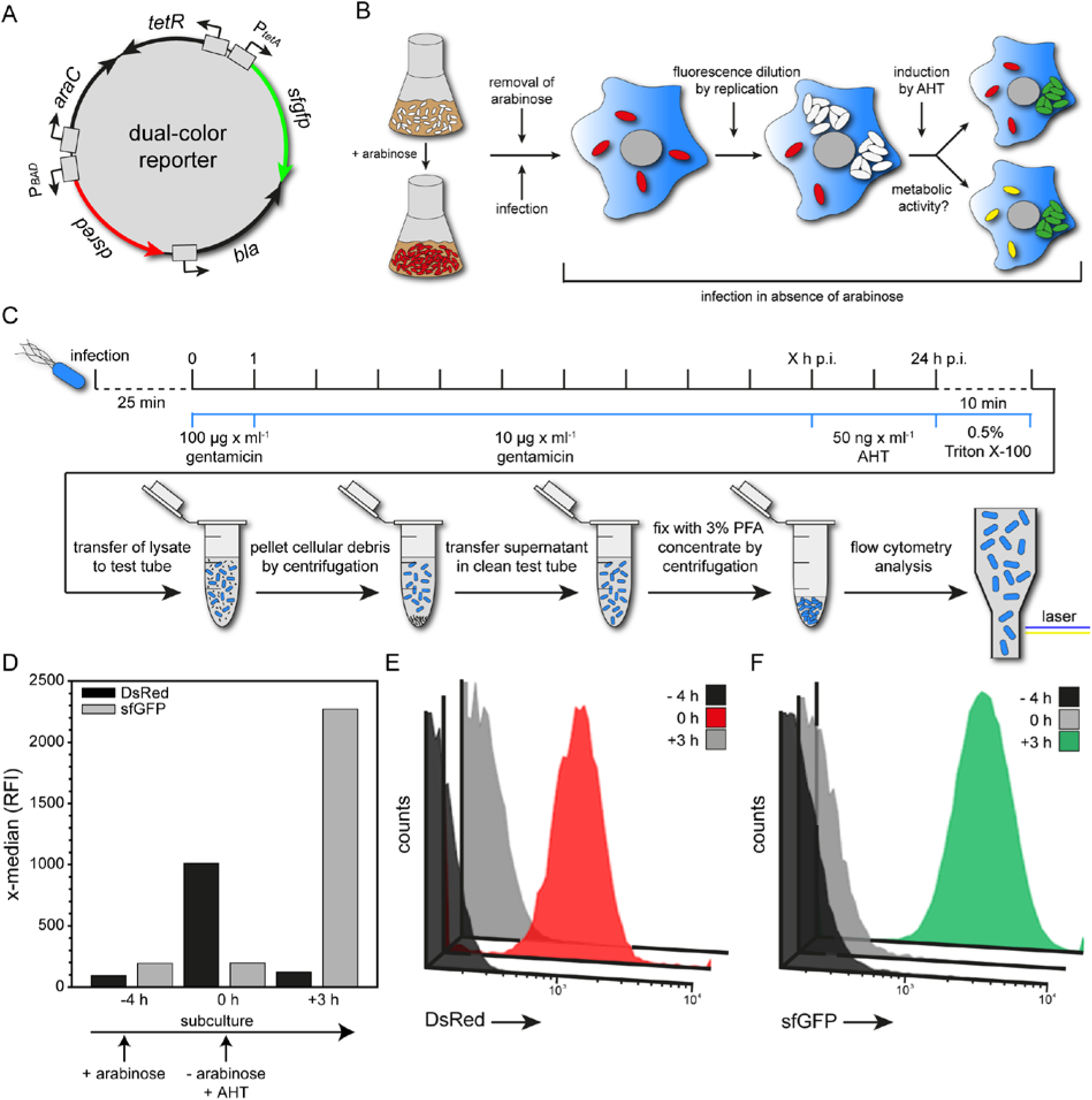
Dual fluorescence reporters for analyses of metabolic activity of intracellular NR STM. A) For the measurement of metabolic activity of NR intracellular STM the EM7 promoter and the *msrA* promoter of the dual fluorescence reporter for *msrA* was replaced by the arabinose-inducible promoter cassette (Guzman et al., 1995), and the tet-ON cassette to be able to artificially induce the expression of *dsred* by arabinose, and *sfgfp* by AHT. B) When adding arabinose to a growing culture of STM in LB medium, bacteria start to synthesize DsRed. Before infection of macrophages, arabinose is removed by centrifugation and washing. Infection is performed without addition of arabinose to the cell culture medium. Replicating intracellular STM lose DsRed via fluorescence dilution. NR intracellular STM keep the DsRed inside the cytoplasm and can be detected as DsRed-positive event after infection. In addition, the metabolic activity can be determined by addition of AHT resulting in synthesis of sfGFP of metabolically active bacteria. C) RAW264.7 macrophages were infected and all non-phagocytosed STM were killed by addition of gentamicin. At indicated time points AHT was added to the cell culture medium. Host cells were lysed 24 h p.i. After removal of cell debris, liberated STM were fixed and subjected to FC. D-F) STM WT harboring double-inducible dual fluorescence reporter was grown in LB medium o/n and diluted in fresh LB medium containing arabinose for further subculture (-4 h). After subculture for 4 h, arabinose was removed by centrifugation and washing (0 h). Subsequently, bacteria were further subcultured for 3 h in fresh LB without arabinose, but containing AHT (+3 h). Samples were taken, fixed and subjected to FC at indicated time points. D) The x-median represents the arabinose-/AHT-induced DsRed/sfGFP intensity of the entire bacterial population. Respective histograms for DsRed intensity (E), and sfGFP intensity (F) are shown.

**Fig. S6:**
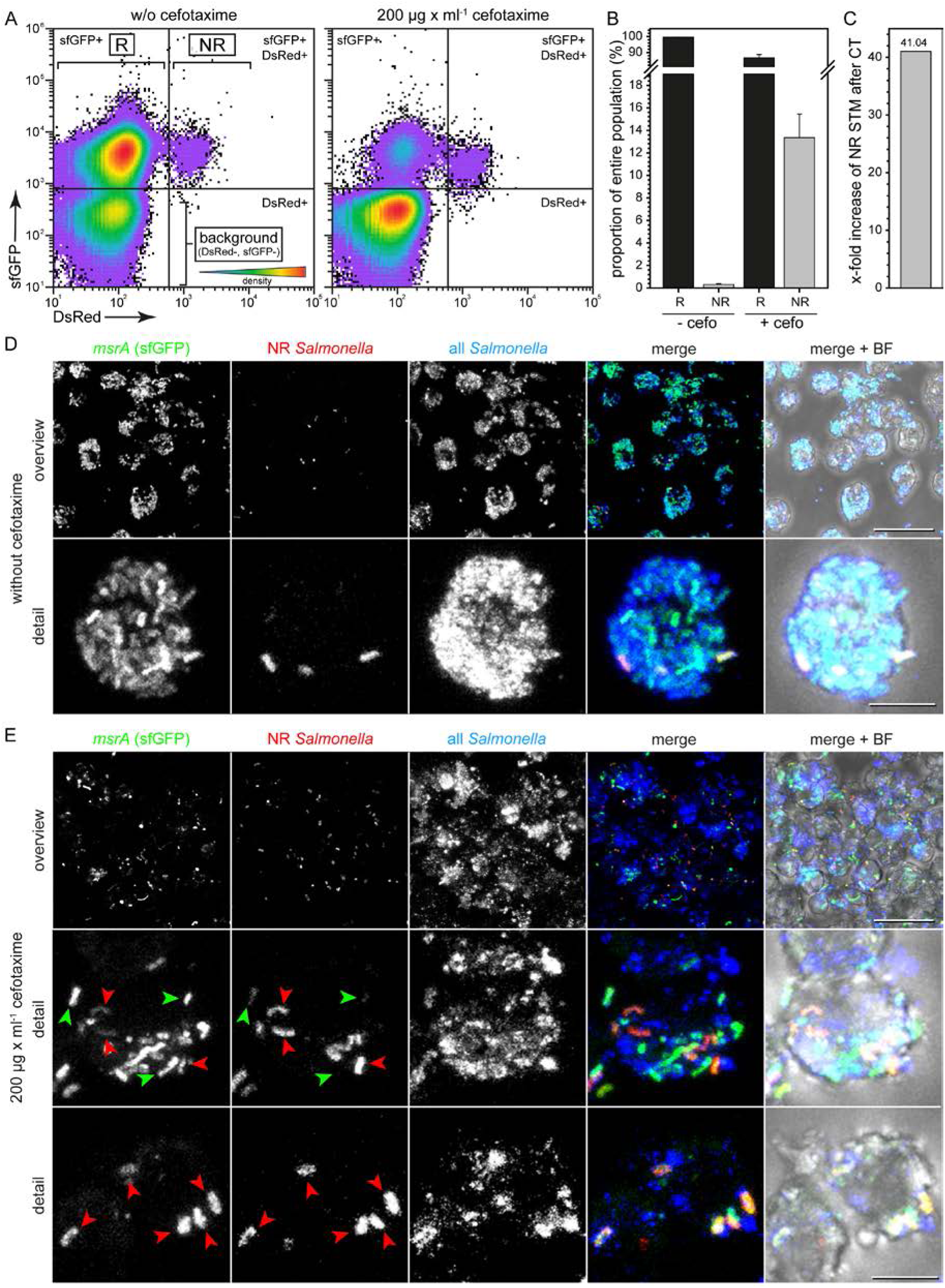
Cefotaxime treatment increases intracellular population of non-replicating STM. WT. STM WT harboring AHT-inducible (NR) dual fluorescence reporter for msrA was grown o/n in LB medium in the presence of AHT. Before infection, AHT was removed. RAW264.7 macrophages were infected, cefotaxime was added 10 h p.i. to the cells if indicated, and cells were lysed for 24 h p.i. FC analyses, or fixed for fluorescence microscopy. Liberated STM were recovered, fixed and subjected to FC. A) Plotting the entire intracellular bacterial population against their AHT-induced DsRed and P*_msrA_*-induced sfGFP intensity shows a high reduction of R STM after antibiotic treatment. B) Quantification of the proportion of R and NR STM within the population with and without antibiotic treatment. C) Calculated x-fold increase of the proportion of NR compared to R STM after cefotaxime treatment (CT). Means and standard deviations of one representative experiment are shown. D, E) Microscopy of intracellular R and NR STM without (D), and with cefotaxime treatment (E). Fixed intracellular STM were immuno-stained against O antigen (blue). Induction of *msrA* is shown in green, NR STM show and R STM do not show red fluorescence. Merge micrographs with and without overlay of brightfield (BF) channels are shown. NR STM are indicated by red arrowheads, R STM are indicated by green arrowheads. Representative cells are shown. Scale bars, 20 µm and 5 µm in overview and detail, respectively.

**Fig. S7:**
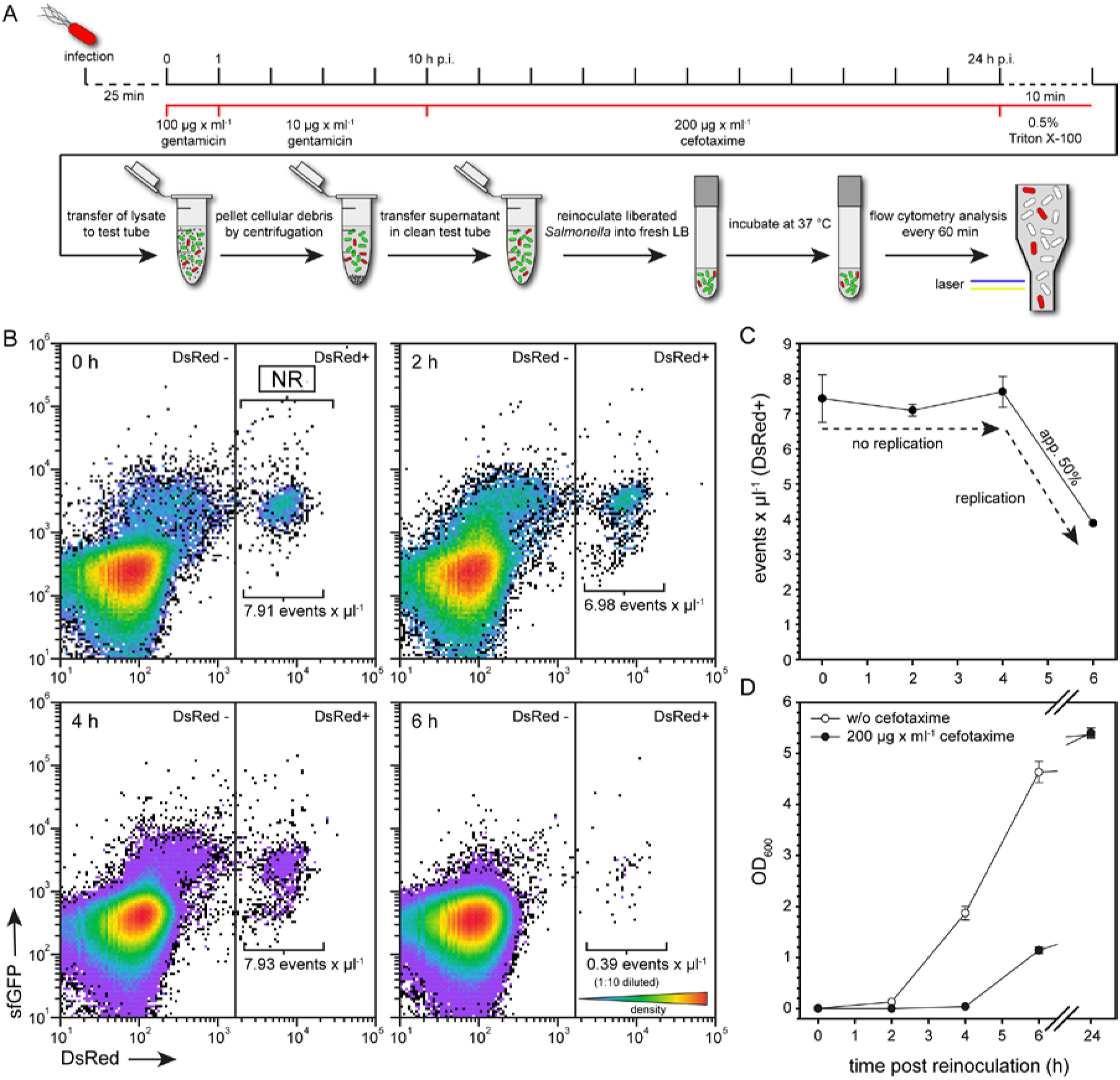
Schematic overview of infection and lysis of host cells, reinoculation into fresh LB, and subsequent cytometric analysis. A) RAW264.7 macrophages were infected using STM harboring various reporters. After killing of all non-phagocytosed STM by addition of gentamicin, 10 h p.i. cell culture medium was replaced by medium containing 200 µg × ml^-1^ cefotaxime. After incubation for further 14 h, host cells were lysed 24 h p.i. After removal of cell debris, liberated STM were reinoculated into fresh LB medium and incubated at 37 °C. At indicated time points samples were taken, fixed and subjected to FC. Quantification of DsRed-positive STM was performed using an Attune NxT cytometer. B) STM WT harboring AHT-inducible (NR) dual fluorescence reporter for *msrA* was grown o/n in LB medium in the presence of AHT. Before infection, AHT was removed. RAW264.7 macrophages were infected, 10 h p.i. cefotaxime was added to the cells and then lysed 24 h p.i. Liberated STM were recovered, reinoculated into fresh LB and incubated at 37 °C. At indicated time points samples were taken, fixed and subjected to FC. C) The relative amount (events × µl^-1^) of DsRed-positive NR STM in the culture was determined. A loss of DsRed-positive events over time indicates that NR STM are growth-competent and start to replicate. D) In addition, the optical density was measured. Means and standard deviations of one representative experiment are shown.

## Notes

### Competing Interest Statement

The authors have declared no competing interest.

